# Inactivation of *lmo0946* (*sif*) induces the SOS response and MGEs mobilization and silences the general stress response and virulence program in *Listeria monocytogenes*

**DOI:** 10.1101/2023.08.28.555070

**Authors:** Magdalena Ładziak, Emilia Prochwicz, Karina Gut, Patrycja Gomza, Karolina Jaworska, Katarzyna Ścibek, Marta Młyńska-Witek, Katarzyna Kadej-Zajączkowska, Eva M.S. Lillebaek, Birgitte H. Kallipolitis, Agata Krawczyk-Balska

## Abstract

Bacteria have evolved numerous regulatory pathways to survive in changing environments. The SOS response is an inducible DNA damage repair system that plays an indispensable role in bacterial adaptation and pathogenesis. Here we report a discovery of the previously uncharacterized protein Lmo0946 as an SOS response interfering factor (Sif) in the human pathogen *Listeria monocytogenes.* Functional genetic studies demonstrated that *sif* is indespensible for normal growth of *L. monocytogenes* in stress-free as well as multi-stress conditions, and *sif* contributes to susceptibility to β-lactam antibiotics, biofilm formation and virulence. Absence of Sif promoted the SOS response and elevated expression of mobilome genes accompanied by mobilization of the A118 prophage and ICELm-1 mobile genetic elements (MGEs). These changes were found to be associated with decreased expression of general stress response genes from the σB regulon as well as virulence genes, including the PrfA regulon. Together, this study uncovers an unexpected role of a previously uncharacterized factor, Sif, as an inhibitor of the SOS response in *L. monocytogenes*.

**SUMMARY:** This study uncovers an unexpected role of a previously uncharacterized factor, Sif, as an inhibitor of the SOS response in *L. monocytogenes*.

## INTRODUCTION

*Listeria monocytogenes* is a Gram-positive foodborne human pathogen that is exquisitely well adapted to survive exposure to severe environmental challenges including high salt concentrations, a wide range of temperatures and extreme pH, and high concentration of β-lactam antibiotics used in the treatment of listeriosis. The versatility of *L. monocytogenes* allows for its adaptation to the natural environment, survival under food processing, succesful infection of host organisms and survival under antibiotic therapy (Hof, 2003; NicAogáin & O’Byrne, 2016). Although infections are not very common, the high mortality rate of listeriosis combined with the widespread dissemination of this organism in the environment makes it a serious public health risk (EFSA, 2022). Regarding this, the primary goals of most research are focused on the detailed understanding of mechanisms which enable survival of *L. monocytogenes* in different environments. Such an understanding could lead to finding the Achilles heel of this pathogen and rational design of new control measures that prevent the survival of *L. monocytogenes* in clinical and non-clinical settings.

A key step in adapting to new stresses is coordination of gene expression with the physiological needs. This is achieved by reprogramming of the transcriptional landscape in response to different physical and chemical signals sensed by *L. monocytogenes*. So far, the best documented role in the response of *L. monocytogenes* to stressful conditions has been established for genes coding for multiple regulatory proteins including, among others, the alternative sigma factor B (σB), the virulence master regulator PrfA, the nutrient-responsive regulator CodY, the RNA chaperone Hfq and two-component signal transduction systems (Bennett et al., 2007; Christiansen et al., 2004; Dorey et al., 2019; Reniere et al., 2016; Williams et al., 2005).

One of the important factors involved in the response of *L. monocytogenes* to multiple stresses is a ferritin-like protein (Fri) belonging to the Dps (DNA-binding proteins from starved cells) family of proteins. The role of Dps-like proteins is to counteract the adverse effects of iron under aerobic conditions (Haikarainen & Papageorgiou, 2010). While Fri of *L. monocytogenes* has no regulatory properties, it contributes to virulence and plays a role in protection against various stresses including acid stress, oxidative stress, iron-starvation, cold- and heat-shock (Dussurget et al., 2005; Milecka et al., 2015; Olsen et al., 2005). Furthermore, inactivation of the *fri* gene was shown to disturb the level of the global regulators catabolite control protein A (CcpA) and anti-σB factor (RsbW), thus indicating the importance of Fri for *L. monocytogenes*’ gene regulatory networks (Dussurget et al., 2005; Milecka et al., 2015). In a previous study, we showed that Fri also is a mediator of β-lactam tolerance and innate resistance to cephalosporins. Interestingly, we observed that *fri* is co-transcribed with the two downstream genes, *lmo0944* and *lmo0945* (Krawczyk-Balska et al., 2012). We hypothesized that genes co-transcribed with *fri* could play an important role in stress adaptation and/or virulence. Therefore, the aim of the present study was to characterize these genes functionally. Firstly, we have undertaken an extended co-transcription analysis revealing that *fri* is transcribed together with four downstream genes, namely *lmo0944*, *lmo0945, lmo0946* and *lhrC5*. Subsequent functional analyses revealed that *lmo0946* of unkown function (conserved in Firmicutes) is indespensible for normal growth of *L. monocytogenes* in a stress-free environement and required under multi-stress conditions. To answer the question why inactivation of *lmo0946* results in severe phenotypic changes of *L. monocytogenes*, we performed comparative RNA-seq analysis of wild type and the mutant strain with inactivated *lmo0946* gene. We found that inactivation of *lmo0946* leads to activation of the SOS response which is a conserved bacterial pathway induced upon various types of DNA damage and regulated by the repressor LexA and the activator RecA. Based on this finding, we propose to rename the *lmo0946* gene to *sif* (<u>S</u>OS interfering factor). Furthermore, inactivation of *sif* leads to downregulation of stress response genes from the σB regulon, virulence genes from the PrfA regulon, and highly upregulated expression of *hfq* encoding an RNA chaperone involved in post-transcriptional regulation of genes in *L. monocytogenes*. Interestingly, inactivation of *sif* resulted in mobilization of two mobile genetic elements (MGEs), namely prophage A118 and ICELm-1. Taken together, our studies led to identification of Sif as an important factor in the physiology of *L. monocytogenes*. Considering the severe phenotypic disorders caused by the inactivation of *sif* in virualy all conditions tested, we propose that Sif represents a good candidate for an Achilles heel of *L. monocytogenes*.

## RESULTS

### Analysis of The Co-transcription of *fri* with Downstream Genes

The study concerning identification of penicillin G-inducible genes of *L. monocytogenes* revealed that *fri* is transcribed together with its downstream genes *lmo0944* and *lmo0945* (Krawczyk-Balska et al., 2012). Further *in silico* analysis of the genomic region comprising the *fri* gene indicated that *lmo0946* and *lhrC5* are located downstream from, and in orientation consistent with, *lmo0945.* This led us to hypothesise that these two genes may also be a part of the *fri* operon (Figure 1a). To evaluate this hypothesis, RT-PCR analysis was performed that verified the co-transcription of *fri*, *lmo0944*, *lmo0945*, *lmo0946* and *lhrC5* genes (Figure 1b). Importantly, in addition to the observed co-transcription event, transcription of *lmo0944*, *lmo0945* and *lhrC5* genes is expected to proceed from their own promoters. To get more detailed information about individual transcripts produced within the ferritin operon and their level of co-transcription, northern blot analysis was performed on total RNA extracted from *L. monocytogenes* EGD-e grown in BHI in exponential phase, with and without penicillin G exposure. The results of the northern blotting, shown in Figure 1c, demonstrate signals corresponding to each gene transcript from the *fri* operon. According to these signals, *fri*, *lmo0944* and *lhrC5* are predominantly transcribed as monocistronic transcripts with size ∼ 500 nt, ∼ 400 nt and ∼ 100 nt, respectively, with the *lhrC5* transcript clearly visible only in presence of penicillin G. For *lmo0945* and *lmo0946*, the signals in the northern blot analysis indicate that these two genes are co-transcribed as no signal with a size corresponding to a monocistronic transcript of any of them is visible. Each of the analyzed genes gave additional bands in the northern blot which may originate from co-transcription with downstream genes and /or processing of the larger co-transcripts. Noteworthy is that signals corresponding to a larger co-transcript with size ∼2300 nt are visible for each of the analyzed genes, providing proof for co-transcription of *fri* with all the genes analyzed. The signal intensity of this co-transcript is much lower than signals corresponding to the monocistronic transcripts of *fri*, *lmo0944* and *lhrC5*, and the bicistronic transcript of *lmo0945* and *lmo0946*, indicating that co-transcription of *fri* with downstream genes does not proceed very effectively in the analyzed conditions of growth. We were wondering if co-transcription level depends on the conditions of growth. To answer this, further northern blot analysis with probes specific for *fri* and *lmo0946* was performed on total RNA extracted from *L. monocytogenes* EGD-e in stationary phase of growth. According to the signals in the northern blot analysis, shown in Figure 1d, *fri* is still predominantly transcribed as the monocistronic transcript with size ∼ 500 nt, however the co-transcript with size ∼2300 nt is clearly visible in stationary phase. In case of *lmo0946*, the intensity of the signal corresponding to this co-transcript is much higher than the signal corresponding to the bicistronic transcript of *lmo0945-lmo0946.* These observations suggest that co-transcription of *fri* with downstream genes occurs more readily in stationary phase cells. Generally, northern blotting confirmed co-transcription of *fri* with *lmo0944*, *lmo0945*, *lmo0946* and *lhrC5* genes. Thus, these genes constitute an operon, named afterwards, the ferritin operon.

**Figure 1.**
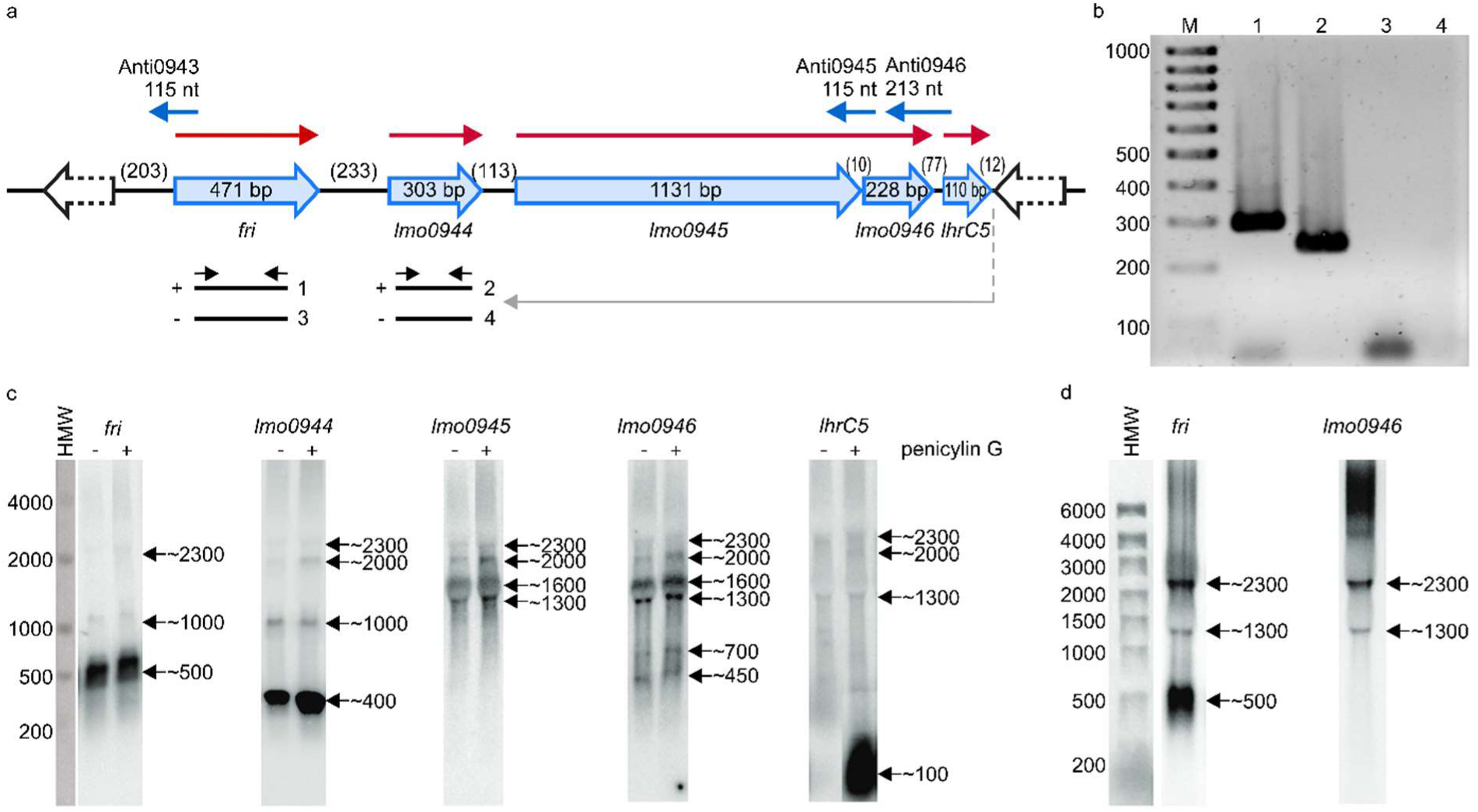
Analysis of co-transcription of *fri* with downstream genes. **(a)** Schematic representation of the genomic region comprising the *fri*, *lmo0944*, *lmo0945*, *lmo0946*, and *lhrC5* genes. Red arrows indicate mRNAs corresponding to the individual genes of the operon and blue arrows represent small antisense transcripts encoded from this genome region. Scheme for co-transcription analysis by RT-PCR is shown below. The template RNA was isolated from exponential-phase cultures of *L. monocytogenes* EGD-e grown in BHI broth at 37°C with 0.09 μg/ml penicillin G. Gray arrow indicates the position of the primer used in RT reaction and black arrows indicate the positions of primers used for PCR. (-) or (+) indicate the expected products amplified on template of RT reactions performed without or with reverse transcriptase, respectively. **(b)** The products obtained in RT-PCR reactions. The numbering of the agarose gel lanes corresponds to the expected products presented in panel (a). The expected size of the amplified fragments of fri and lmo0944 was 288 bp and 212 bp, respectively. A 100-bp ladder (lane M) is shown as a size marker. **(c)** Northern blot analysis of *fri*, *lmo0944*, *lmo0945*, *lmo0946*, and *lhrC5* in exponential-phase of *L. monocytogenes* EGD-e growth. Samples were taken from EGD-e wild type mid-exponential cultures exposed to 1 hour of penicillin G stress (+) as well as from non-stressed cultures (-). Northern blots were probed for *fri* mRNA, *lmo0944* mRNA, *lmo0945* mRNA, *lmo0946* mRNA, and LhrC5. Numbers on the right side of each panel indicate the estimated lengths of the transcripts in nucleotides. **(d)** Northern blot analysis of *fri* and *lmo0946* in stationary-phase of *L. monocytogenes* EGD-e growth. Samples were taken from EGD-e wild type overnight culture. Northern blots were probed for *fri* mRNA and *lmo0946* mRNA. Numbers on the right side of each panel indicate the estimated lengths of the transcripts in nucleotides.

### Inactivation of *lmo0946* impairs the Growth under Different Conditions

To investigate the relevance of the ferritin operon in the physiology of *L. monocytogenes*, each gene of the operon was subjected to inactivation. In case of *lmo0944* and *lmo0945*, in frame deletion mutants were constructed. Due to the presence of genes coding for antisense RNAs Anti0943, Anti0945 and Anti0946 within the operon (Figure 1a), single point mutations were designed for inactivation of *fri*, *lmo0946* and *lhrC5* to minimize the risk of undesirable changes in the expression of these asRNAs. In case of *fri* and *lmo0946* the nucleotide substitutions were introduced to create nonsense mutations (named *fri*\* and *lmo0946*\*), and in case of *lhrC5*, nucleotide substitutions in the -10 promoter region were introduced to prevent initiation of transcription (named *lhrC5*\*). Moreover, due to high sequence similarity between LhrC5 and six other sRNAs from the LhrC family, the functional analysis LhrC5 may be difficult or impossible when the other LhrCs remain functional. To resolve this problem, the genes encoding for these six LhrCs were deleted in the background of the wild-type and *lhrC5* mutant strains. First, the growth rates of the mutants and the parent strain in BHI broth at 37°C were compared. Interestingly, in these stress-free conditions the inactivation of all the studied genes but *lmo0946* had no effect on rate of growth (Figure 2).

**Figure 2.**
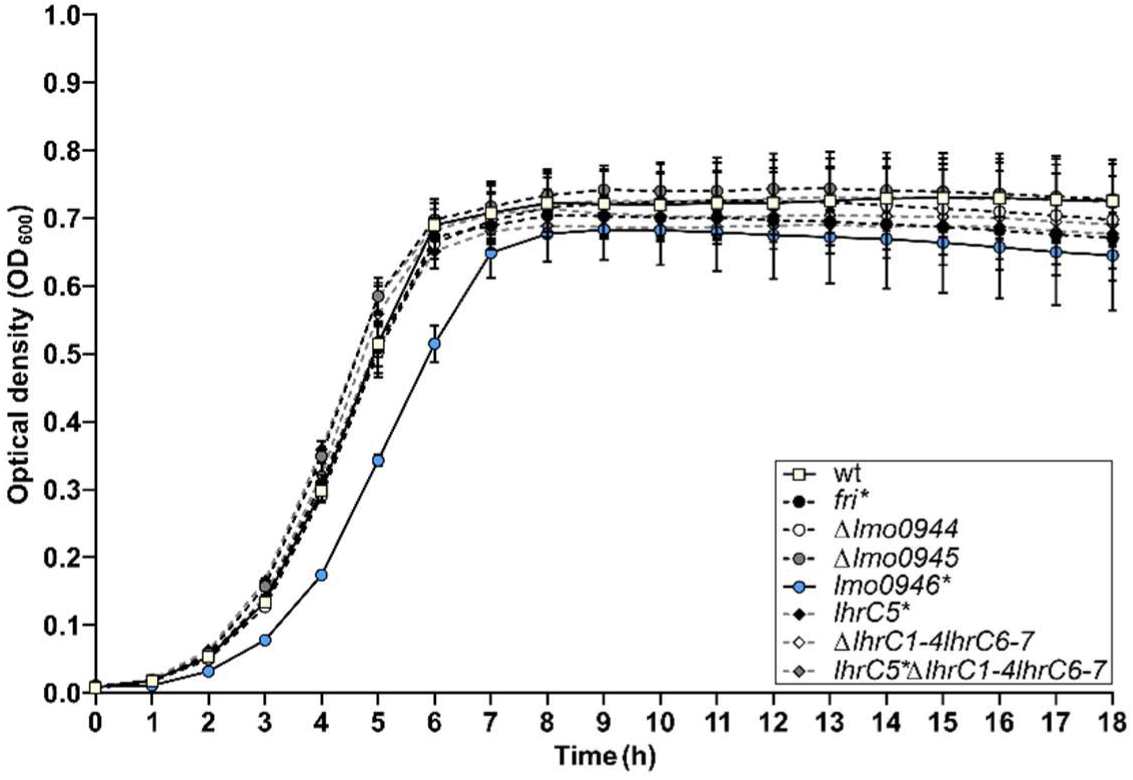
Growth of wild-type *L. monocytogenes* EGD-e and the ferritin operon mutants in stress-free conditions. BHI broth was inoculated with an overnight culture of each strain (1:1000) and incubated with shaking at 37°C. Cell growth was measured spectrophotometrically by determining the OD_600_. The mean values from three independent experiments are plotted and the error bars represent the standard deviation.

The observed growth impairment of the *lmo0946\** mutant strain prompted us to focus on assessing the effects of inactivation of this gene. Prior to detailed analysis, complementation of *lmo0946* was performed in the background of the mutant strain. The wild-type, *lmo0946\** mutant and complementated strain (*lmo0946*\*-*lmo0946*) were compared with respect to their abilities to grow in stress-free conditions. The growth of the *lmo0946\** mutant was clearly impaired relative to the wild type and *lmo0946*-lmo0946* strains (Figure 3a, Supplementary Table S1). Subsequent analysis of ability of the *lmo0946\** mutant to grown under various stress conditions, including the presence of subinhibitory concentrations of ethanol and penicillin G, acid and alkaline environments, or high temperature, revealed that Lmo0946 is inecessary for normal growth of *L. monocytogenes* under various environmental conditions (Figure 3b-3f, Supplementary Table S1).

**Figure 3.**
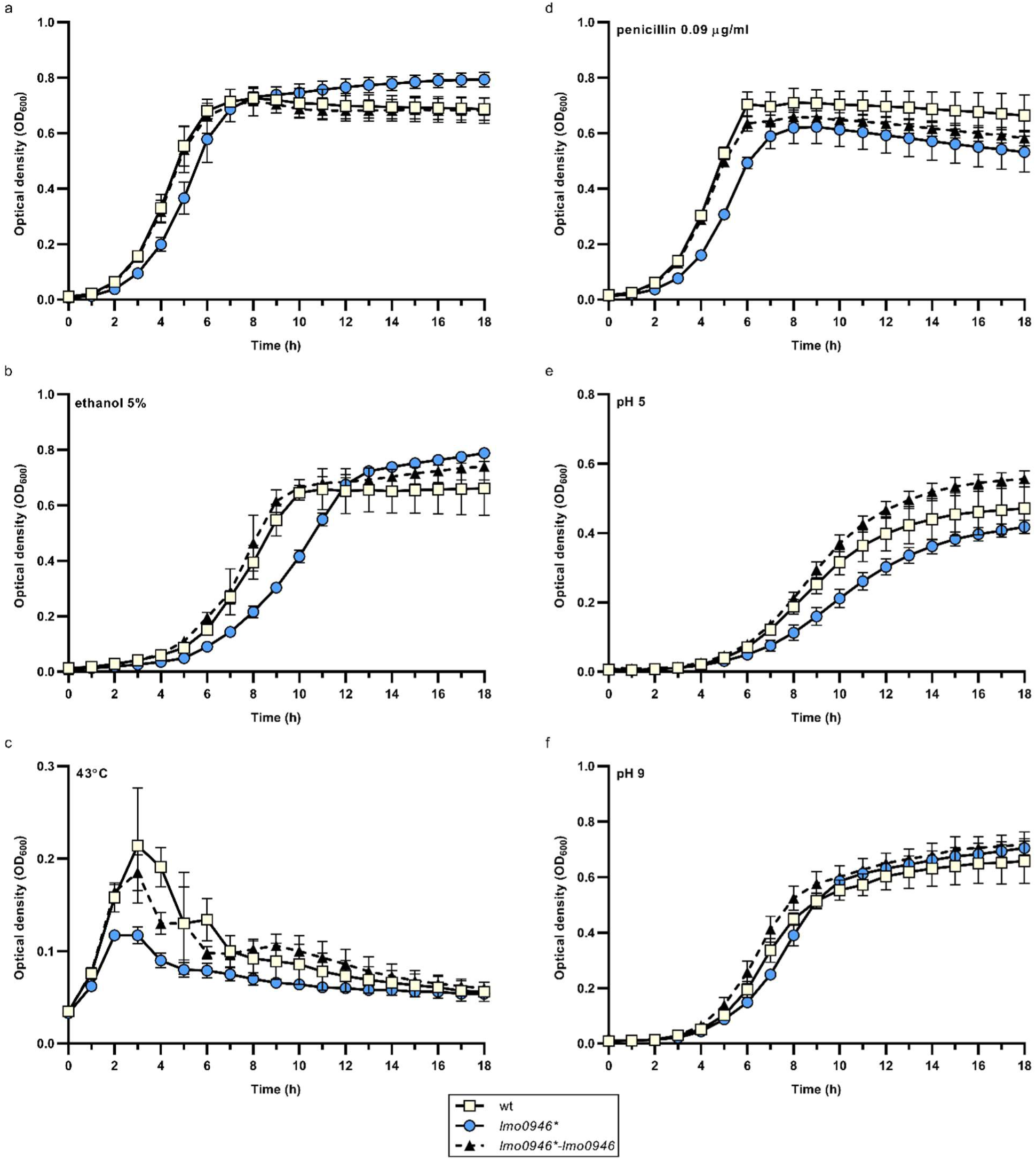
Inactivation of *lmo0946* impairs growth of *L. monocytogenes* under different conditions. Growth of wild-type *L. monocytogenes* EGD, *lmo0946*\* mutant and the complementated strain *lmo0946*-lmo0946* in **(a)** BHI broth, **(b)** BHI broth suplemented with 5 % ethanol, **(c)** BHI broth at 43°C, **(d)** BHI broth suplemented with 0.09 µg/ml penicillin G, **(e)** BHI broth pH 5 **(f)** and BHI broth pH 9. BHI broth was inoculated with an overnight culture of each strain (1:1000) and incubated with shaking at 37°C except high temperature stress **(c)**. Cell growth was measured spectrophotometrically by determining the OD_600_. The mean values from three independent experiments are plotted and the error bars represent the standard deviation. Calculated generation times are shown in Supplementary Table S1.

### Inactivation of *lmo0946* reduces the ability of *L. monocytogenes* to form biofilm

In order to study the effect of *lmo0946* inactivation on biofilm formed by *L. monocytogenes*, the sessile biomass of the studied strains was stained by crystal violet. The biofilm formation assay revealed significant differences between the *lmo0946\** mutant, and wild-type or complemented strains (Figure 4). The amount of sessile biomass for the *lmo0946\** mutant was significantly diminished up to 72 h of sessile growth compared to the wild-type and complemented strains (Figure 4). These results indicate that inactivation of *lmo0946* reduces the ability of *L. monocytogenes* to create biofilm.

**Figure 4.**
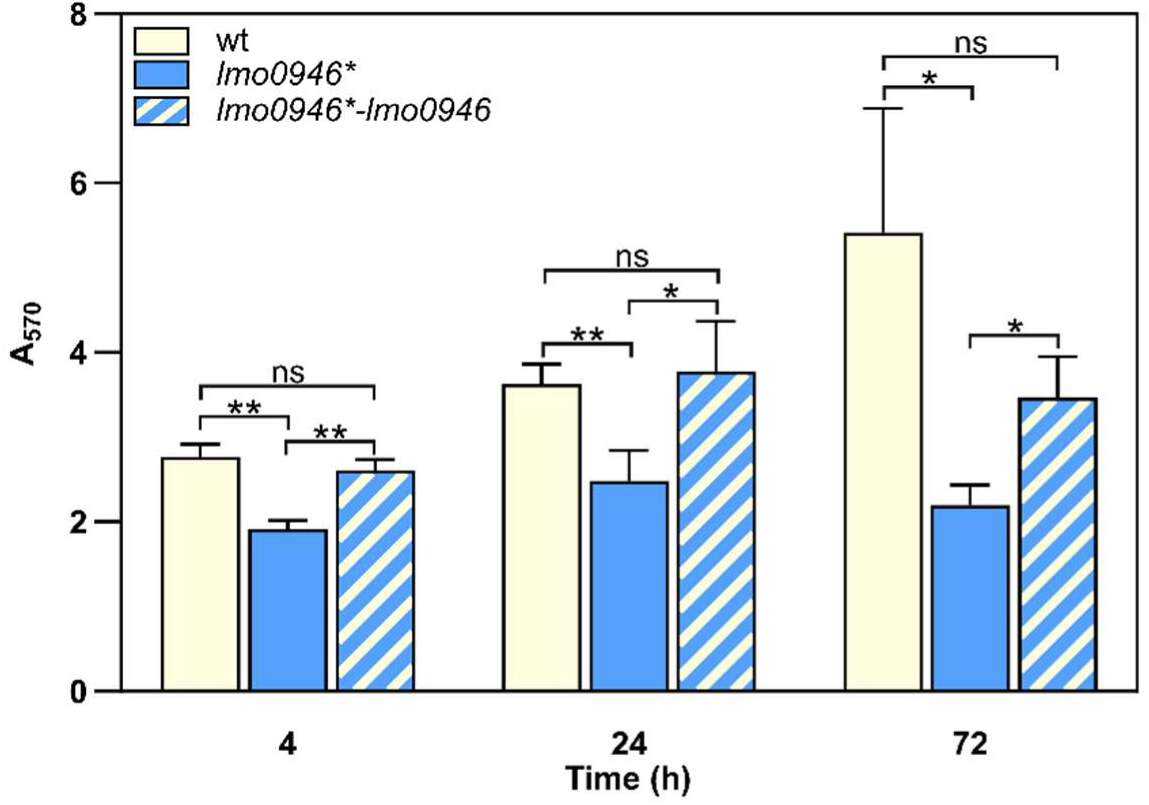
Inactivation of *lmo0946* reduces biofilm formation by *L. monocytogenes*. Sessile development of wild-type *L. monocytogenes* EGD-e, *lmo0946\** mutant and the complementated strain *lmo0946*-lmo0946* are assayed with the crystal violet method. The data represent the means and SD of three biological repeats. The asterisks indicate a significant differences (*P < 0.05, **P < 0.01).

### Lmo0946 contributes to the resistance of *L. monocytogenes* to cephalosporins and tolerance to penicillin G

To investigate whether inactivation of *lmo0946* affects the susceptibility of *L. monocytogenes* to antibiotics, the parent and mutant strains were subjected to antibiotic disk assays. This assay revealed that the wild-type and *lmo0946*\* strains displayed similar levels of resistance to various antibiotics (ampicillin, penicillin G, vancomycin, aztreonam, meropenem, tetracycline, rifampicin, gentamicin, trimethoprim, and ciprofloxacin), however significantly greater zones of growth inhibition were observed for the mutant with cefuroxime and cefoxitin. The MICs of these specific cephalosporin antibiotics were then determined for *L. monocytogenes* EGD-e and the *lmo0946*\* mutant. In confirmation of the antibiotic disk assay result, the MIC of cefuroxime for wild-type and *lmo0946*\* was 8 μg/ml and 4 μg/ml, respectively, whereas the MIC of cefoxitin for wild-type and *lmo0946*\* was 32 μg/ml and 16 μg/ml, respectively. Thus, inactivation of the *lmo0946* gene caused a 2-fold increase in the sensitivity of *L. monocytogenes* to these cephalosporins. Next, tolerance was examined by testing the ability of the strains to survive in concentrations of penicillin G over 100 fold higher than the MIC value which is 0.12 μg/ml. The tolerance assay revealed that the survival of *lmo0946*\* was significantly impaired since reduced numbers of viable cells were recovered for the mutant relative to the wild-type strain after exposure to high concentration of penicillin G (Figure 5). In summary, our results indicate that Lmo0946 contributes to the resistance to cephalosporins and tolerance to penicillin G of *L. monocytogenes*.

**Figure 5.**
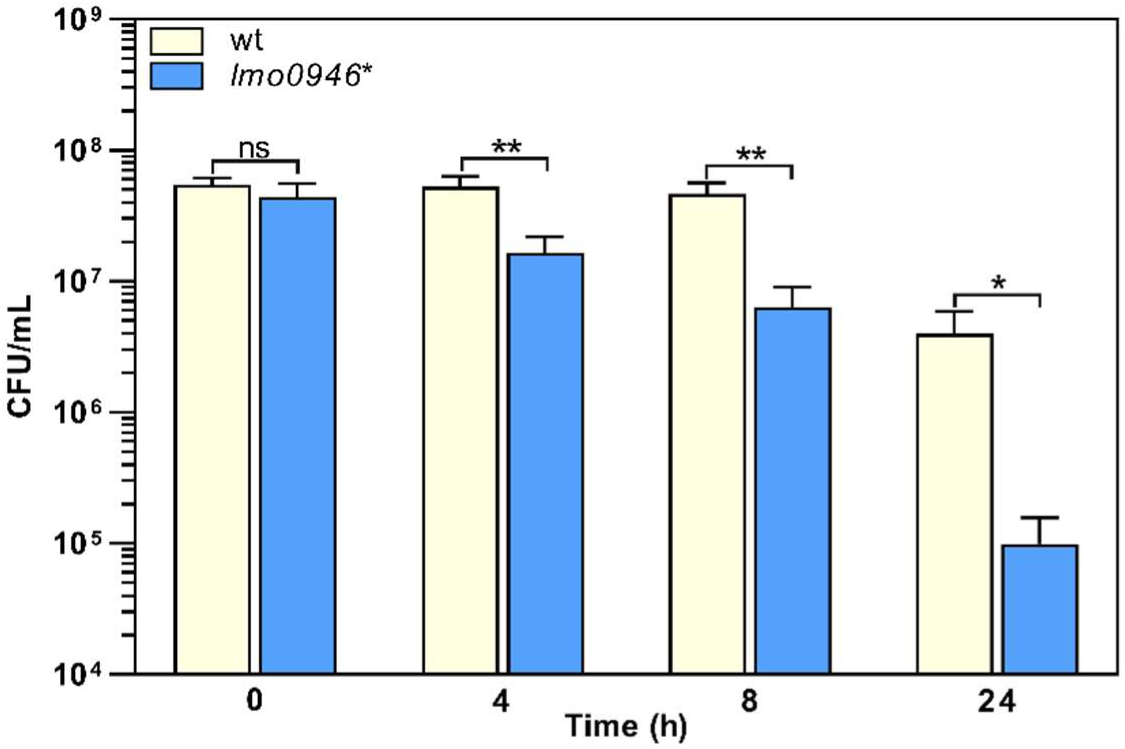
Inactivation of *lmo0946* impairs tolerance of *L. monocytogenes* to penicillin G. BHI broth supplemented with 32 μg/ml penicillin G was inoculated with a mid-exponential culture of wild-type *L. monocytogenes* EGD-e (wt) and *lmo0946\** mutant strains and incubated with shaking at 37°C. Viable cell counts were measured on agar plates following serial dilutions of culture samples. The mean values from three independent experiments are plotted and the error bars indicate standard deviations. The asterisks indicate a significant differences (*P < 0.05, **P < 0.01).

#### Inactivation of *lmo0946* impairs the virulence of *L. monocytogenes*

To determine whether Lmo0946 is important for the virulence of *L. monocytogenes*, *in vitro* infection studies were performed using the *lmo0946\** mutant and wild-type *L. monocytogenes* EGD-e. The infection assay showed that the *lmo0946\** mutant was significantly impaired in the ability to proliferate intracellularly in the murine macrophage cell line P388D1 relative to the wild-type strain (Figure 6a). Subsequently, the mouse infection model was used to assess the virulence properties of the *lmo0946\** mutant relative to that of wild-type. For the mutant, survival and/or growth in the liver and spleen was significantly reduced at day 3 post-infection when compared with wild-type (Figure 6b). These results indicate that Lmo0946 contributes to the virulence of *L. monocytogenes*.

**Figure 6.**
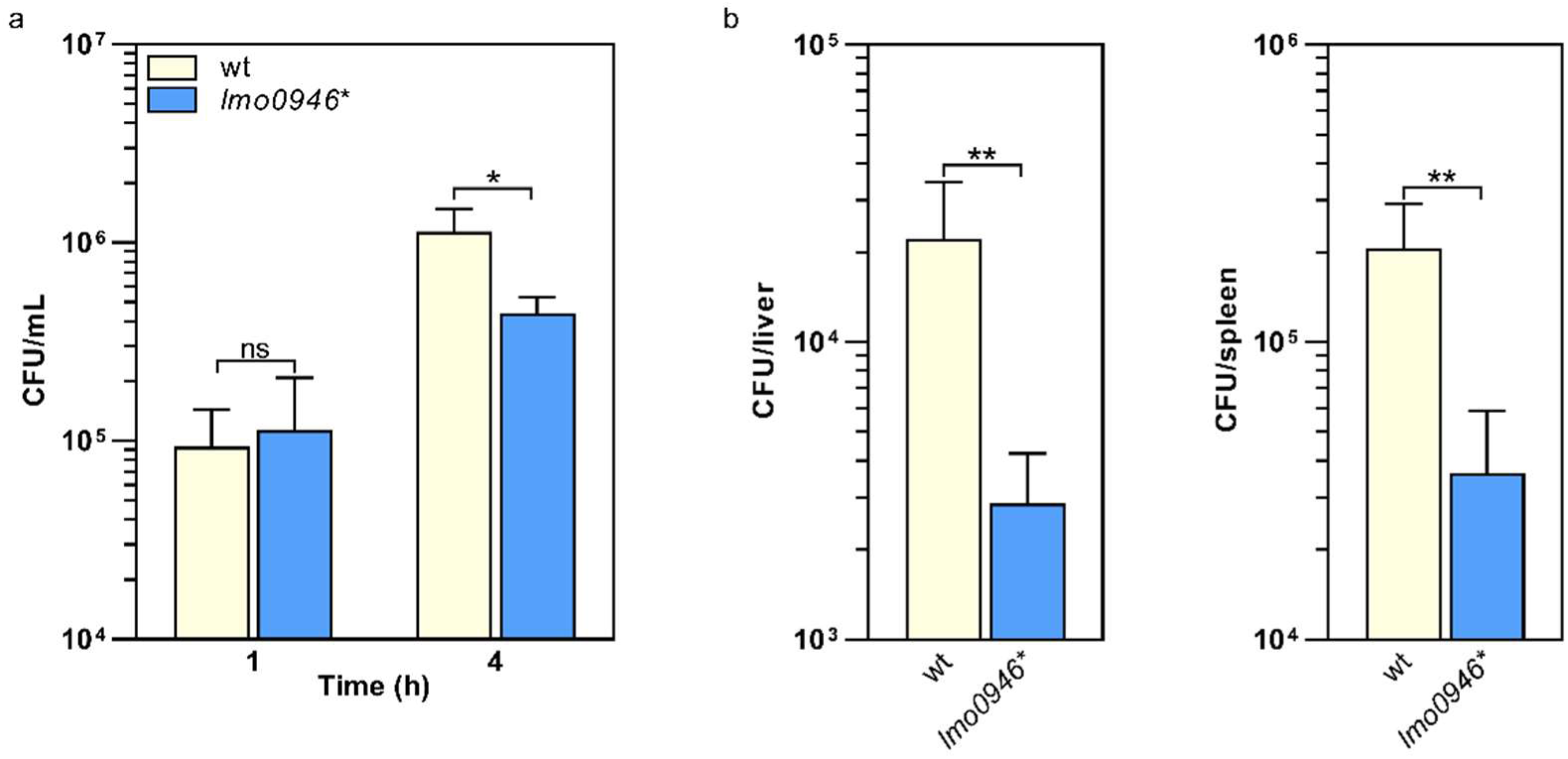
Inactivation of *lmo0946* impairs virulence of *L. monocytogenes*. **(a)** The effect of *lmo0946* inactivation on the intracellular replication of *L. monocytogenes* in the murine macrophage cell line P388D1. Cell monolayers were infected with the wild-type *L. monocytogenes* EGD-e (wt) and *lmo0946\** mutant strains with an MOI of 10 in 24-well plates. Bacterial CFU counts were measured on agar plates following lysis of the infected cells at the indicated time of post-infection. The mean values from three independent experiments are plotted and the error bars indicate standard deviations (*P < 0.05). **(b)** Mice infection with wild-type *L. monocytogenes* EGD-e and *lmo0946\** mutant strains. Bacterial load in BALB/c mice organs were determined following intravenous inoculation with 2000 CFU of *L. monocytogenes* EGD-e as well as *lmo0946\** mutant. On day 3 after infection, the numbers of viable bacteria in livers and spleens of five animals per group were determined. Error bars indicate standard deviations. The asterisks indicate a significant differences (*P < 0.05, **P < 0.01).

### Inactivation of *lmo0946* Causes Global Changes of Genes Expression Including Upregulation of the Mobilome and Downregulation of Metabolism

*In silico* analysis of Lmo0946 indicates that it is a small (75 aa) conserved bacterial protein of unknown function, and the function of its homologs, found mainly in *Bacilli* and *Clostridia* classes of the *Firmicutes* phylum (Supplementary Figure S1), has not been established yet. To further elucidate why inactivation of *lmo0946* results in severe phenotypic changes of *L. monocytogenes*, a transcriptomic analysis of *L. monocytogenes lmo0946\** mutant and the parental strain EGD-e was performed using RNA-seq during exponential phase of growth at 37°C in BHI. The RNA-seq analysis revealed that inactivation of *lmo0946* caused 589 genes to be differentially expressed when compared to the wild-type strain using an adjusted p-value < 0.01, corresponding to approximately 20 % of protein encoding genes and 10 % of the sRNAs (Figure 7a and Supplementary Table S2). The relative abundance of each COG category in the set of differentially transcribed genes was further determined. A set of 320 genes showed higher transcript levels while 269 showed lower transcript levels in the *lmo0946\** mutant relative to the wild-type strain (Figure 7b and Supplementary Table S3). COGs analysis revealed that genes involved in carbohydrate transport and metabolism, energy production and conversion, and lipid transport and metabolism were highly represented in the downregulated set of genes, indicating that inactivation of *lmo0946* results in a general slowdown of metabolism. The analysis also showed that genes related to nucleotide transport and metabolism, replication, recombination and repair as well as the mobilome were highly represented among the upregulated genes, suggesting that inactivation of *lmo0946* leads to disorders of processes related to DNA metabolism and replication.

**Figure 7.**
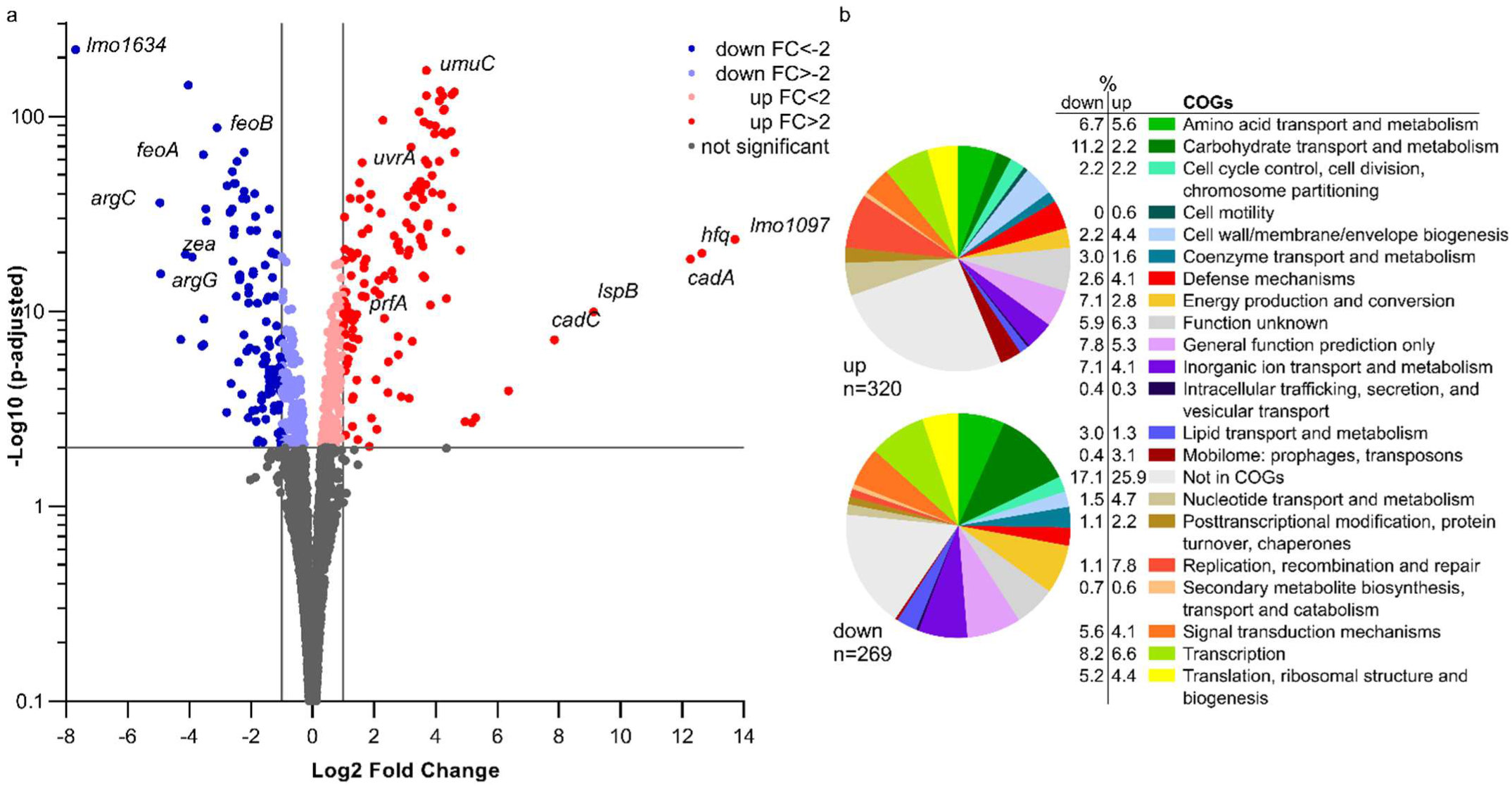
RNA-seq analysis of genes differentially transcribed by *L. monocytogenes lmo0946\** mutant grown to exponential phase in BHI medium. **(a)** Volcano plots displaying differentially expressed genes in *L. monocytogenes lmo0946*\* mutant compared to the wild-type EGD-e. The negative log10 of an adjusted p-value is plotted on the Y-axis, and the log2FC is plotted on the X-axis. The horizontal line represents an adjusted p-value > 0.01 cutoff. The vertical lines represent a – 1.0> log2FC > 1.0 cutoffs. **(b)** Relative abundance of categories of genes differentially expressed by *L. monocytogenes lmo0946*\* mutant comparatively to the wild-type EGD-e corresponding to categories in the COGs database. Upper pie chart, genes upregulated in the mutant compared with wild-type; Lower pie chart, genes downregulated in the mutant compared with wild-type.

To better understand the causes of global changes in gene expression resulting from *lmo0946* inactivation we further analyzed the genes that were at least fourfold up- or downregulated in the background of *lmo0946\** (corresponding to 83 and 45 genes, respectively) (Supplementary Table S4). The most up-regulated genes with more than 20-fold change were *lmo1097* (encoding an integrase), *hfq* (encoding an RNA chaperone), genes involved in cadmium resistance, and few genes coding for hypothetical proteins (Supplementary Table S4 and Figure 7a). Going through the most numerous parts of the up-regulated genes, it was noticeable that most of them are phage-related genes. These genes accounted for over 50% of the upregulated genes and most of them, though not all, were genes of the A118 prophage which is one of the four mobile genetic elements (MGEs) found in the *L. monocytogenes EGD-e* strain. The three other MGEs are the monocin (or *lma*) locus, the integrative and conjugative element ICELm1 (TN916-like) and IS3-1 (Kuenne et al., 2013).

Given the high frequency of prophage genes in the set of up-regulated genes, the expression of all MGE genes in the *lmo0946*\* strain was further analyzed. This analysis showed that genes of three of the four MGEs, i.e., prophage A118, monocin locus and ICELm-1, are highly expressed in the *lmo0946*\* mutant (Supplementary Table S5).

The monocin locus, a cryptic prophage region, is conserved in all *L. monocytogenes* lineages and includes the *lmaDCBA* operon with an important role in virulence of *L. monocytogenes* (Hain et al., 2012). The expression of all genes from the monocin locus is up-regulated in the *lmo0946*\* mutant, and for most of them more than 10-fold up-regulation is observed. ICELm1 is an MGE specific for *L. monocytogenes* EGD-e. It consists of 19 genes, including an integrase-encoding gene (*lmo1097*), cadmium resistance genes *lmo1100* (*cadA*)*, lmo1101* (*lspB*) and *lmo1102* (*cadC*), a gene coding for a fibrinogen-binding protein with an LPXTG domain (*lmo1115*) and Tn916-like genes of unknown function (Kuenne et al., 2013; Parsons et al., 2018; Pombinho et al., 2017). In case of this MGE, eight genes were found to be upregulated in *lmo0946*\* including the afromentioned genes with known function (Supplementary Table S5). Interestingly, genes belonging to ICELm1 showed over 30-fold up-regulation of expression in the *lmo0946*\* mutant strain, and this was the highest increase in expression recorded in these studies. An analysis of the A118 prophage showed that the expression of 50 of 62 prophage genes is up-regulated in the *lmo0946*\* strain with more than 4-fold up-regulation observed for 45 of these genes (Supplementary Table S5). Interestingly, the upregulated genes belong to virtually all functional modules of prophage A118 including genes encoding phage lysis proteins holin and lysin. In summary, these data indicate that inactivation of *lmo0946* led to upregulation of a large portion of the mobilome of *L. monocytogenes*.

The most highly down-regulated gene in the *lmo0946\** mutant was *lmo1634*. It encodes the Listeria adhesion protein (LAP), a putative bifunctional acetaldehyde-CoA/alcohol dehydrogenase involved in pyruvate metabolism. Three pyruvate-formate lyase encoding genes were found to be downregulated as well (*pflB*, *pflA* and *pflC*). Additionally, genes involved in arginine biosynthesis (*argC, argG, argH*), ferrous iron transport (*feoA* and *feoB*), and amino acid transport and metabolism (*arpJ* and *gadD*) were found to be highly down-regulated in the *lmo0946\** mutant relative to the wild-type strain (Supplementary Table S4 and Figure 7a). Collectively, these findings suggest that inactivation of *lmo0946* led to a decrease in energy production and conversion as well as descreased transport and metabolism of amino-acids and other cell compounds. Interestingly, high level of downregulation was also noticed for *lmo2686* encoding the RNA binding protein Zea which was shown to specifically interact with phage RNAs of *L. monocytogenes* (Pagliuso et al., 2019). This observation further supports a putative link between *lmo0946* and the mobilome of *L. monocytogenes*.

### Inactivation of *lmo0946* Triggers a Variety of Stress Responses in *L. monocytogenes*, Including the Sos Response

Out of 128 genes with at least a 4-fold change in expression level, 68 were found to be allocated into known regulons, and some of them are under control of more than one regulator (Supplementary Table S4). For these genes the largest overlap was observed with the CodY regulon. The transcriptional regulator CodY responds to different nutritional and environmental stresses and generally controls adaptive responses in conditions that limit bacterial growth (Bennett et al., 2007). We noticed that 30 of the highly upregulated genes, coding for prophage A118 proteins, were under positive control of CodY. In addition, seven of the most downregulated genes were under negative control of CodY. Among them were *arg* genes coding for proteins engaged in the biosynthesis of arginine, and genes involved in the transport and metabolism of amino acids, carbohydrates, and inorganic ion.

In addition to the CodY regulon, five genes belonging to LexA/RecA regulon were upregulated at least fourfold in *lmo0946\** mutant strain (Supplementary Table S4). The LexA/RecA regulon covers genes involved in the SOS response that is induced upon various types of DNA damage (van der Veen et al., 2010). This observation led us to further analyze the effect of inactivation of *lmo0946* on level of expression of the SOS regulon. The analysis revealed that nearly all of the 29 genes of LexA/RecA regulon are upregulated in the mutant strain, except from the σB-dependent genes *bilEA* and *bilEB* involved in osmoprotection (Sue et al., 2003), and *lmo2266* with unknown function (Supplementary Table S6). Notably, the expression levels of genes from the SOS response noticed for the *lmo0946\** mutant strain is comparable to the levels expressed in wild-type *L. monocytogenes* EGD-e after treatment with the SOS-inducing agent mitomycin C (van der Veen et al., 2010). Among the most highly induced genes were those encoding the cell division inhibitor YneA (l*mo1303*), the translesion DNA polymerases DinB (*lmo1975*) and UmuDC (*lmo2675* and *lmo2676*) (Supplementary Table S6). These findings clearly demonstrate that inactivation of *lmo0946* triggers the expression of SOS response genes known to promote the DNA repair in DNA-damaged cells. Finally, four genes belonging to the VirR regulon were upregulated at least fourfold in the *lmo0946\** mutant strain. VirR is an important regulator of genes involved in bacterial surface components modifications (Mandin et al., 2005). Most of the VirR-dependent upregulated genes code for proteins involved in amino acid transport and metabolism (Supplementary Table S4).

### Inactivation of *lmo0946* Leads to Downregulation of Virulence Genes and Modulation of Expression of σB and CodY Regulons

An analysis of 45 highly downregulated genes revealed that eight of them belong to the PrfA regulon (Supplementary Table S4). PrfA is the master virulence regulator of *L. monocytogenes* (Reniere et al., 2016). Further analysis of the effect of inactivation of *lmo0946* on the PrfA regulon revealed that expression of 29 from 73 genes was changed with an adjusted P value <0.01 in the *lmo0946\** mutant strain and further 12 genes with an adjusted P value <0.05 (Supplementary Table S7). Interestingly, all differentially expressed genes from the PrfA regulon, except *prfA and lmo0641*, are downregulated in the *lmo0946\** mutant, even though nearly all of them correspond to genes under positive control by PrfA (Supplementary Table S7). Furthermore, decreased level of expression was observed for genes preceded by a PrfA box and belonging to the core set of the PrfA regulon, like *hly*, *inlA* and *inlB* (Milohanic et al., 2003). Additionally, the most highly downregulated gene, *lmo1634*, encodes the LAP protein that promotes bacterial translocation across the intestinal epithelial barrier (Drolia et al., 2018). Furthermore, among the highly downregulated genes were *arpJ* and *pplA* coding for factors playing important roles at different stages of intracellular infection of *L. monocytogenes* but not belonging to PrfA regulon (Supplementary Table S4; Reniere et al., 2016). In addition to the PrfA regulon, five genes belonging to the σB regulon were downregulated at least fourfold in the *lmo0946\** mutant strain (Supplementary Table S4). The alternative sigma factor σB positively regulates genes involved in the response of *L. monocytogenes* to multiple stress conditions and has also been implicated in virulence (Dorey et al., 2019). σB-dependent downregulated genes encode proteins with unknown function, except for *lmo2067* (*bsh)* coding for the bile salt hydrolase with an important role in intestinal phase of infection (Begley et al., 2005). As observed for the PrfA regulon, σB positively regulates the genes found to be downregulated in the *lmo0946\** mutant (Supplementary Table S4). The only gene positively regulated by σB, and highly upregulated in the *lmo0946\** strain, is *hfq* coding for an RNA chaperone involved in post-transcriptional control of gene expression in *L. monocytogenes*. Notably, transcription of *hfq* was shown to proceed from a σB-dependent promoter (Christiansen et al., 2004). These data suggest that modulation of expression of the σB regulon occurs in the *lmo0946\** strain. This assumption is supported by a 2-fold decrease in expression of the *rsbV* gene coding for the anti-anti-σB factor (Supplementary Table S2). As *rsbV* plays an important role in a signaling pathway controlling σB activity (Dorey et al., 2019), this observation prompted us to gain insight into the expression of the *rsb* genes coding for components of the stressosome as well as the σB signaling cascade. The *rsb* genes are encoded within the *sigB* operon and are responsible for sensing and subsequent transmission of stress signals to activate the σB factor (Dorey et al., 2019). The analysis revealed that in addition to *rsbV* also *rsbW* encoding an anti-σB factor is slightly but significantly decreased in the *lmo0946\** mutant strain (Supplementary Table S8). This observation suggests that the σB signaling cascade is affected by the *lmo0946* mutation. Among the highly downregulated genes were also three genes downregulated by Fur, which is a Fe^2+^ dependent repressor of genes involved in iron/heme uptake and utilization. These genes encode a ferrous iron transport system (*feoA* and *feoB*) and an element of a transport system involved in cytochrome biosynthesis (*cydC*) (Lechowicz & Krawczyk-Balska, 2015).

It should be noted that as many as 16 out of 25 highly downregulated genes belonging to known regulons are regulated in *lmo0946\** in a manner opposite to regulation by the appropriate regulators (Supplementary Table S4). Besides the genes from the PrfA and σB regulons described above, seven genes positively regulated by CodY also belong to this group. These observations indicate that inactivation of *lmo0946\** result in global modulation of expression of genes belonging to different regulons involved in stress response and virulence of *L. monocytogenes*.

### Validation of transcriptomic changes observed in the *lmo0946\** mutant strain by RT-qPCR analysis

To ensure that the performed RNA-seq analysis reflects the transcriptional changes in the *lmo0946*\* mutant strain, the RNA-seq data was subsequently validated by RT-qPCR analyses of 12 genes (including up-, non- and down-regulated genes). The obtained results indicated very good correlation between RNA-seq and RT-qPCR results (R^2^=0.945; (Supplementary Figure S2). Subsequently, to verify whether the transcriptomic changes observed in the *lmo0946*\* mutant strain are the result of inactivation of *lmo0946*, the level of expression of selected genes in wild-type *L. monocytogenes EGD-e,* the *lmo0946\** mutant and the complemented strain was examined by RT-qPCR. The analysis of genes from the SOS regulon and mobile genetic elements revealed increased level of expression in the *lmo0946*\* mutant (Figure 8a). Upon complementation, the expression of these genes was restored to that observed in the wild-type strain, except for the *lmo1097* gene belonging to ICELm1 MGE (Figure 8a). Next, the expression of virulence genes was examined. Increased expression of *prfA* and decreased expression of *hly* was observed in the *lmo0946*\* mutant, and upon complementation the expression of these genes was restored to that observed in the wild-type strain (Figure 8b). Finally, the expression of genes with potent regulatory roles was studied. Increased expression of *hfq* and decreased expression of *zea* was observed in the *lmo0946*\* mutant, but upon complementation the expression of these genes was not restored to that observed in the wild-type strain. In case of *rsbV* and *sigB*, expression of these genes is decreased in the *lmo0946\** mutant strain, however only in case of *rsbV* the difference in expresion between wild-type and mutant strain was significantly important (P=0.04), while for *sigB* the difference was just above the cut-off of statistical significance (P=0.08). Furthermore, the expression of *rsbV* was not restored upon compementation to the level observed in wild-type EGD-e strain (Figure 8c). In conclusion, the results of the RT-qPCR analysis confirmed the changes in gene expression in the mutant strain observed in the RNA-seq analysis. Furthermore, gene expression was restored in the complemented strain for genes involved in the SOS response and virulence as well as two MGEs i.e., prophage A118 and monocin locus. However, the expression of genes involved in RNA binding and the σB response as well as ICELm1 MGE was not restored upon complementation.

**Figure 8.**
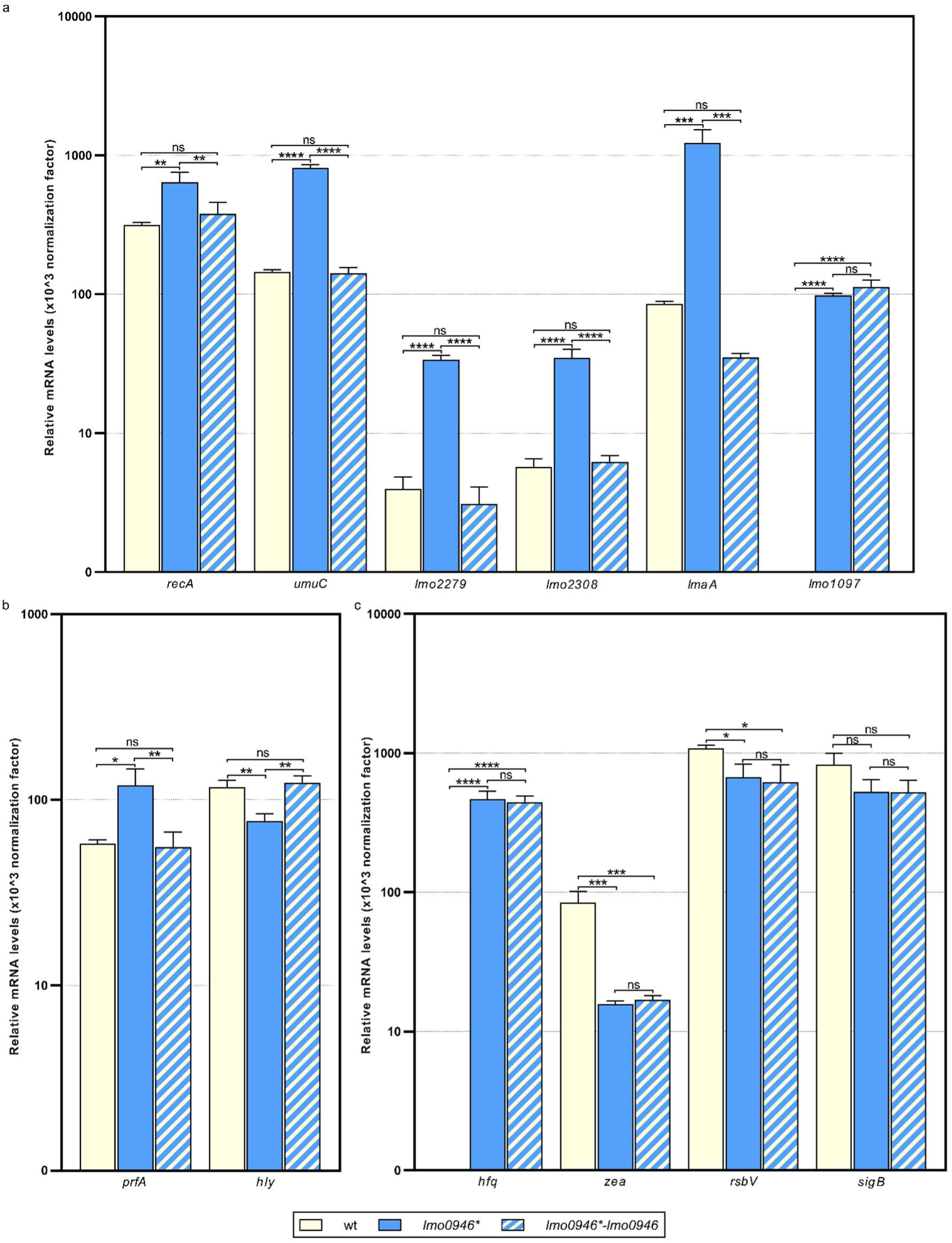
Impact of *lmo0946* inactivation and complementation on expression of selected SOS regulon, MGE, virulence and regulatory genes. The expression was determined in exponential growth phase in BHI for wild-type EGD-e, lmo0946* mutant strain and complemented strain lmo0946*-lmo0946 by RT-qPCR. Relative transcripts levels of **(a)** recA, umuC, lmo2279, lmo2308, lmaA and lmo1097, **(b)** prfA and hly, and **(c)** hfq, zea, rsbV and sigB were normalized to the amount of rpoB gene. The results shown are the average of three biological replicates, error bars indicate standard deviations. The asterisks indicate a significant differences (*P < 0.05, **P < 0.01, ***P < 0.001, ****P < 0.0001).

#### Inactivation of *lmo0946* Results in excision of prophage A118 without inducing lysogeny

The prophage A118 is integrated within the *comK* gene of *L. monocytogenes EGD-e* and while it was not extensively studied, it exhibits high similarity to Φ10403S prophage of *L. monocytogenes* 10403S which was shown to excise without switching from lysogeny to the lytic pathway in the process named active lysogeny (Pasechnek et al., 2020; Rabinovich et al., 2012). Based on the highly increased expression of prophage A118 genes in the *lmo0946\** mutant strain, we hypothesized that excision of the phage could occur. To address this hypothesis, phage DNA excision and extra-chromosomal replication in the wild-type, *lmo0946\** mutant and complemented strain were evaluated by PCR amplification of the intact *comK* gene and a fragment of circularized phage genome, which are formed only upon the phageA118 excision (Figure 9a). This analysis revealed the presence of a PCR product corresponding to a fragment containing the phage genome integration site in the *lmo0946\** mutant strain and lack of this product in the wild-type and complemented strain. Furthermore, an intact *comK* gene was not amplified in the wild-type and complemented strain, while the amplification proceeded efficiently on DNA isolated from the *lmo0946\** mutant strain (Figure 9b). The sequence analysis of PCR products representing intact *comK* and circular phage-DNA confirmed a precise excision of the phage A118 genome in the *lmo0946\** mutant strain which leaves an in-frame coding sequence of the *comK* gene containing *attB* site and a reconstituted phage attachment *attP* site (Figure 9c). These observations indicate that in the *lmo0946\** mutant strain precise excision of prophage A118 is induced and it is reversed upon complementation.

**Figure 9.**
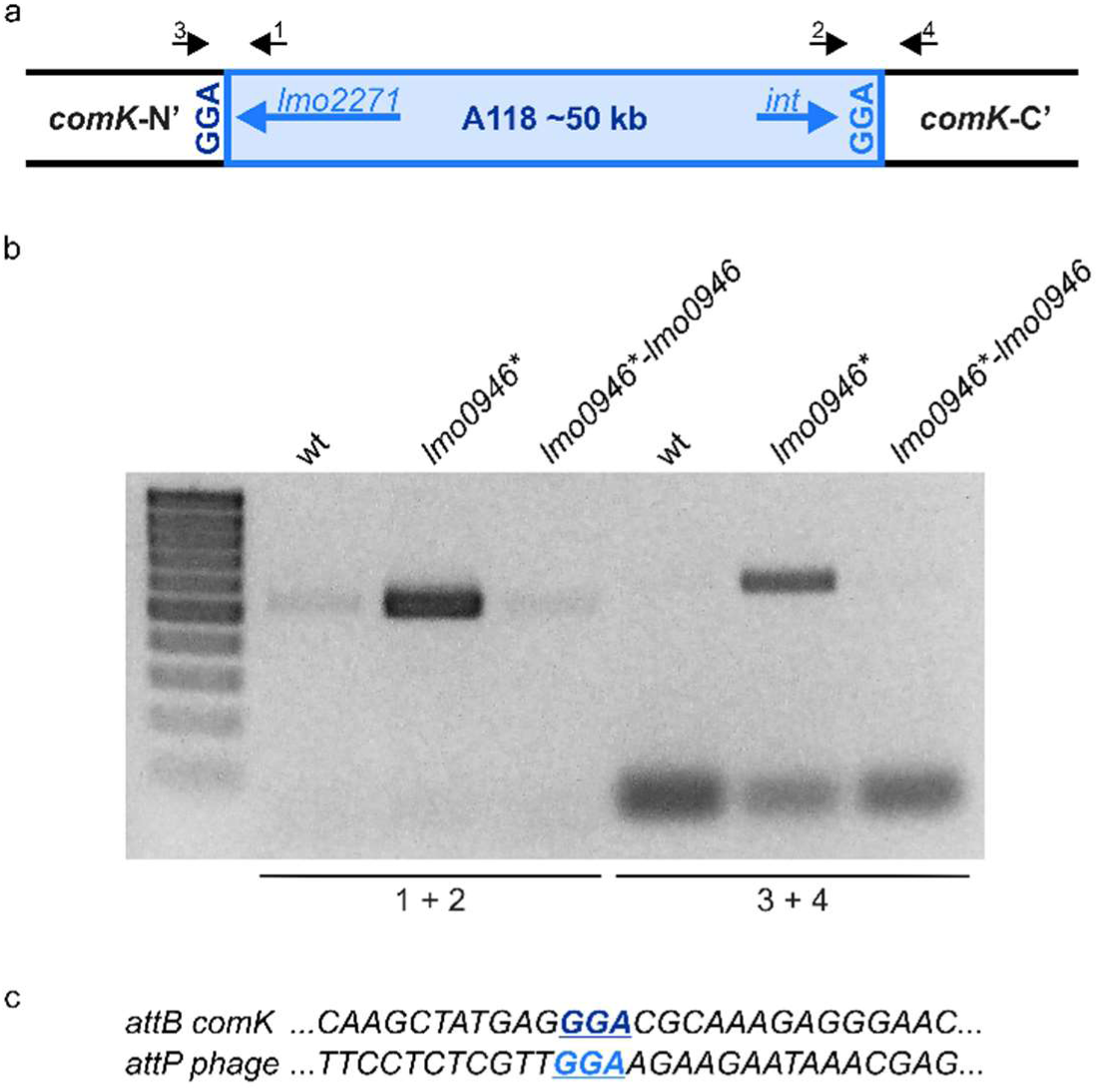
The Prophage A118 Is Excised in *lmo0946*\* mutant strain. **(a)** Schematic representation of the genomic region containing the split *comK* gene with integrated the A118 prophage. Black arrows depict primers used to characterize the *comK*-phage genomic region (primers 1-4). **(b)** PCR analysis of *comK*-phage genomic region in wild-type, *lmo0946*\* mutant and the complemented strain *lmo0946*-lmo0946* grown to exponential (mid-log) phase in BHI medium. **(c)** DNA sequences of the *comK attB* site and the phage *attP* site.

We further wondered whether the observed phage excision in *lmo0946\** mutant strain is followed by induction of the lytic cycle. To answer this question, infective virion production was monitored in the wild-type, *lmo0946\** mutant and complemented strains by using a plaque forming assay. The infective phages production was not detected in any of the strains tested (data not shown). Therefore, prophage A118 does not go into the lytic cycle, even though excision of the prophage occurs in the *lmo0946\** mutant strain. Likewise, PCR analysis was performed to detect possible excision of cryptic prophage monocin locus as a high level of expression of this MGE in *lmo0946*\* mutant was also observed. However, no PCR products were obtained in this analysis indicating that excision of monocin locus in *lmo0946\** mutant does not occur (data not shown).

### Inactivation of *lmo0946* Results in replication of ICELm1 in circular form and increased resistance to cadmium

The integrative and conjugative element ICELm1 is specific for the EGD-e strain and accordingly, it is absent in other widely used *L. monocytogenes* strains. So far, nothing is known about the biological function or mobilization of ICELm1 (Bécavin et al., 2014; Kuenne et al., 2013). As the genes from ICELm1 were found among the most higly upregulated genes in the *lmo0946\** mutant strain, we decided to investigate the possible excision of this MGE. To address this hypothesis, ICELm1 DNA excision and extra-chromosomal replication in the wild-type, *lmo0946\** mutant and complemented strain were evaluated by PCR amplification of a fragment covering the distal *guaA* and proximal *lmo1116* genes as well as a fragment of circularized ICELm1, which both are formed only upon the ICELm1 excision (Figure 10a). This analysis revealed the presence of a PCR product corresponding to the fragment containing the ICELm1 integration site in the *lmo0946\** mutant and complemented strains, however this product was not detected in the wild-type strain. Surprisingly, the fragment covering distal *guaA* and proximal *lmo1116* genes was not amplified in any of the tested strains suggesting the presence of ICELm1 in the genome of the studied strains. This assumption was evaluated by an additional PCR amplification of an intergenic region of *guaA* and *lmo1097* which revealed the presence of a PCR product corresponding to intact intergenic region of *guaA* and *lmo1097* in all studied strains (Figure 10b). The sequence analysis of PCR products representing circular ICELm1 confirmed a precise replication of the circular form of ICELm1 in the *lmo0946\** mutant strain with preserved duplicated attachment *attR* site and resulted in the arising of a truncated form of sRNA *rli38*, named *rli38-1* (Figure 10c). These observations indicate that while the presence of ICELm1 is preserved in all the strains, a precise replication of the circular form of ICELm1 is induced in the *lmo0946\** mutant strain, and it is not reversed upon complementation.

**Figure 10.**
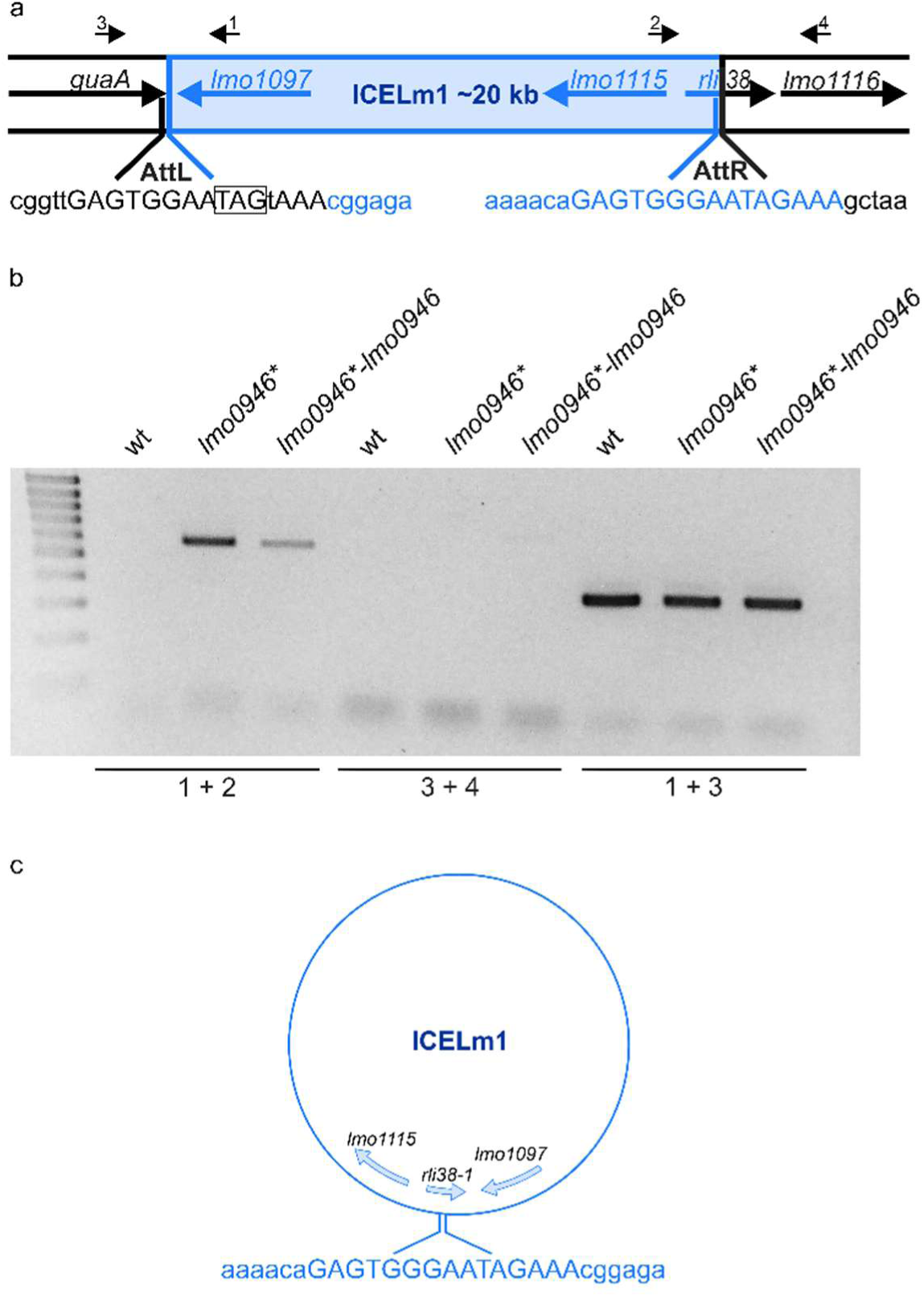
The ICELm1 is replicated and present in circular form in *lmo0946*\* mutant strain. **(a)** Schematic representation of the genomic region containing ICELm1. Black arrows depict primers used to characterize the ICELm1 genomic region (primers 1-4). The *aatL* and *attR* sequences are given in uppercase and stop codon of guaA is boxed. **(b)** PCR analysis of ICELm1 genomic region in wild-type, lmo0946* mutant and the complemented strain grown to exponential (mid-log) phase in BHI medium. **(c)** Schema of circular form of ICELm1 with given sequence of *attR* site.

As the genes involved in cadmium resistance are encoded within the ICELm1 (Kuenne et al., 2013) we further wondered whether the observed replication of the circular form of ICELm1 in the *lmo0946\** mutant and complemented strain led to an increase in cadmium resistance of these strains. To answer this question, the susceptibility of the wild-type, *lmo0946*\* mutant and complemented strains to cadmium was tested using an agar and broth dilution assay. This assay revealed that the MIC of cadmium for EGD-e was 200 μg/ml while the *lmo0946*\* and complemented strains were extremely resistant to cadmium. The precise determination of the MIC values for these strains was impossible due to their ability to grow in the presence of cadmium to the limit of its solubility, which was 1000 μg/ml. These observations indicate that replication of the circular form of ICELm1 in the *lmo0946\** mutant and complemented strain is accompanied by high increase of cadmium resistance.

### Lmo0946 protein does not bind to DNA of promoters of selected genes

Frequently, pronunced changes in bacterial transcriptomics and physiology are observed upon inactivation of regulatory genes. Considering that inactivation of Lmo0946 resulted in major changes in gene expression, we wondered whether Lmo0946 could be a DNA-binding transcriptional regulator. To investigate this idea, we performed electrophoretic mobility shift assays (EMSAs) using native polyacrylamide gels to detect binding of the purified Lmo0946-His_6_ protein to promoter regions of selected genes with clearly changed expression in the mutant strain and important roles in physiology or stress response of *L. monocytogenes*. EMSA analysis showed that Lmo0946-His_6_ lacks the ability to bind DNA fragments corresponding to the promoter regions of *recA*, *prfA*, *lap*, *lmo1097*, *hfq*, *cadA* and *argC* (Supplementary Figure S3). These results indicate that Lmo0946 itself has no DNA binding properties and changes in expression of the analyzed genes in the mutant strain seem not to result from direct regulation of transcription by Lmo0946.

## DISCUSSION

The ferritin-like protein plays an important role in protection against multiple stresses and virulence of *L. monocytogenes* (Dussurget et al., 2005; Milecka et al., 2015; Olsen et al., 2005). We previously showed that the *fri* gene is co-transcribed together with its downstream genes *lmo0944* and *lmo0945* (Krawczyk-Balska et al., 2012). As genes co-transcribed with *fri* could play an important role in stress adaptation and/or virulence, this observation prompted us to extend the co-transcription analysis and functional analyses of genes transcribed together with *fri*. The RT-qPCR revealed that *fri* is co-transcribed with four downstream genes, namely *lmo0944*, *lmo0945, lmo0946* and *lhrC5*, the last of which encodes for an sRNA from the LhrC family known for its role in the stress response of *L. monocytogenes* (Sievers et al., 2014, 2015). As the result from the co-transcription analysis contradicts previous studies showing that Fri is expressed from a monocistronic mRNA both in stress-free conditions and under heat- or cold-shock (Hébraud & Guzzo, 2000), we further analyzed the transcripts of the *fri* operon by northern blot. The analysis showed that the larger co-transcript, corresponding to the mRNA of *fri* and downstream genes, is clearly visible in the stationary phase of growth, however it is barely detectable during growth under stress-free conditions and under penicilin G pressure. These results indicate that the transcriptional terminator of *fri* is regulated in a condition-specific manner that results in generation of full-length mRNA at higher levels in the stationary phase than in other conditions. Therefore, the co-transcription of *fri* with downstream genes could be easily missed in earlier studies. Subsequently, all the genes from the ferritin operon were subjected to functional analysis. The inactivation of *fri, lmo0944, lmo0946 and lhrC5* had no effect on the growth of *L. monocytogenes* in stress-free conditions which in the case of *fri* and *lhrC5* confirms the results of previous studies (Dussurget et al., 2005; Olsen et al., 2005; Sievers et al., 2014), however inactivation of *lmo0946* led to markedly slower growth and reduced number of bacteria. This surprising observation prompted us to focus further research on *lmo0946*. Our analyses revealed that inactivation of *lmo0946* caused slowdown of growth of *L. monocytogenes* in presence of subinhibitory concentrations of ethanol and penicillin G, in acid and alkaline environments, and high temperature. The *lmo0946\** mutant strain also exhibited reduced tolerance to penicillin G, a phenotype that so far is known to be determined by the σB factor and Fri protein (Begley et al., 2006; Krawczyk-Balska et al., 2012). Furthermore, the *lmo0946*\* mutant strain showed increased sensitivity to some cephalosporins – antibiotics to which *L. monocytogenes* shows high innate resistance (Krawczyk-Balska & Markiewicz, 2016). The experiments examining the effect of *lmo0946* inactivation on sessile growth revealed an important role of *lmo0946* in biofilm formation by *L. monocytogenes* as well. Finally, the mutant with inactivated *lmo0946* exhibited attenuated virulence in *in vitro* and *in vivo* models of infection. Overall, the results of the phenotypic analysis indicate that inactivation of *lmo0946* results in severe disorders in the physiology of *L. monocytogenes* in virtually all conditions tested. To further explore the role of Lmo0946 in *L. monocytogenes*, a transcriptomic analysis of the *lmo0946*\* mutant and the parental strain was performed in stress-free conditions, as we assumed that the functional impairment of the mutant strain is related to the transcriptomic changes already observed in stress-free conditions. Our experiments revealed that the response of *L. monocytogenes* to *lmo0946* inactivation is multifaceted. Firstly, *L. monocytogenes* responds to *lmo0946* inactivation by induction of the SOS response which generally is initiated by accumulation of single-stranded DNA in the cell. This situation results in the activation of RecA, which in turn stimulates autocatalytic cleavage of represor LexA and, finally, the induction of the SOS response (Maslowska et al., 2019). Notably, RecA-mediated activation of the SOS response factor YneA, responsible for cell division arrest in *L. monocytogenes* (van der Veen et al., 2010), explains the observed growth retardation of the *lmo0946\** mutant in different conditions of growth. The SOS response can be induced by a plethora of factors, including physical and chemical DNA damaging-agents present in the environment, and DNA-damaging byproducts and intermediates of cellular metabolism, such as reactive oxygen species (ROS). Since the SOS response in the mutant is induced during growth without harmful factors, it must be related to the changes in cellular processes that occur because of the mutation. The induction of the SOS response observed in the *lmo0946\** mutant is not likely caused by induced production of ROS since the transcriptome analysis did not reveal an induction of *kat* (encoding catalase) and *sod* (encoding superoxide dismutase) known to work together in detoxification of ROS (Dallmier & Martin, 1988). Furthermore, these genes were recently reported to be highly co-induced with the SOS response in *L. monocytogenes* exposed to heme stress causing oxidative damage (Dos Santos et al., 2018). This conclusion is further supported by the observation that genes belonging to the PerR regulon dealing with peroxide stress in *L. monocytogenes* (Rea et al., 2005) are lacking among the genes showing greatly upregulated expression in the mutant strain (Supplementary Table S4). Worth of note is that different stimuli can indirectly generate the SOS-inducing signal by activation of endogenous DNA damage mechanisms (Aertsen & Michiels, 2006). Given this, we speculated that Lmo0946 may be a transcriptional regulator of genes with important roles in the physiology or stress response of *L. monocytogenes* and its regulatory activity in turn could lead to activation of the SOS response. However, the results of EMSA analyses led us to reject this hypothesis. Therefore, the specific mechanism underlying the link between Lmo0946 inactiation and activation of the SOS response remains unclear at the current stage of research. Although the present study does not explain how the SOS system is activated in the *lmo0946*\* mutant, the complementation studies show that this effect is clearly dependent on the *lmo0946* gene. Based on this finding, we propose to rename *lmo0946* gene to *sif* (<u>S</u>OS interfering factor).

Generally, one of the well-known effects of the induction of the SOS response is activation of mobile genetic elements including endogenous prophages (Little & Mount, 1982). Accordingly, we observed phage A118 mobilization in the *sif* mutant strain. The temperate A118 prophage is specific to *L. monocytogenes* serovar 1/2 strains and belongs to the Siphoviridae family of double-stranded DNA bacterial viruses (Zink & Loessner, 1992). This bacteriophage was shown to reproduce by both lytic and lysogenic cycles, in the latter the phage’s genome is integrated at a specific attachment site located within the *comK* gene (Loessner et al., 2000). Surprisingly, we observed that phage A118 mobilization is not accompanied by switching from lysogeny to the lytic pathway in a *sif* mutant strain of *L. monocytogenes* EGD-e. This result is consistent with previous reports on hardly induction of the lytic cycle of prophage A118 in the EGD-e strain exposed to UV irradiation or mitomycin C; well known factors inducing the SOS response (Pasechnek et al., 2020). Notably, a similar prophage to A118, named Φ10403S, is integrated like A118 in the *comK* gene of another widely used *L. monocytogenes* laboratory strain, 10403S. Upon UV irradiation or mitomycin C treatment the Φ10403S phage enters the lytic cycle, while the excision without production of progeny virions is specifically induced during intracellular growth to promote bacterial escape from macrophage phagosomes. This process, named active lysogeny, led to modulation of the virulence of *L. monocytogenes* 10403S in the course of infection (Pasechnek et al., 2020; Rabinovich et al., 2012). In the light of these data, we assume that the interplay between *comK*-phages and their *L. monocytogenes* hosts is strain dependent and the active lysogeny triggered by the SOS response in the *sif* mutant may be a specific response of the *L. monocytogenes* EGD-e strain. The active lysogeny in *L. monocytogenes* 10403S is ensured by negative control of the expression of the Φ10403S phage’s late genes, including the *lysin* and *holin* genes that mediate bacterial lysis (Pasechnek et al., 2020). In contrary, we observed a high level of transcription of A118 phage late genes, including *holin* and *lysin*. This result indicates that the control of active lysogeny of phage A118 in strain EGD-e is different form that of the Φ10403S phage in strain 10403S. The high level of transcription of *holin* and *lysin* of A118 phage suggests a post-transcriptional control of active lysogeny in the *sif* mutant of the EGD-e strain. This hypothesis may be supported by the observed changed transcription in the *sif* mutant strain of two genes, namely *hfq* and *zea*, with known and putative roles, respectively in post-transcriptional regulation of gene expression in *L. monocytogenes.* The Hfq of *L. monocytogenes* is an RNA chaperone involved in post-transcriptional regulation of gene expression (Nielsen et al., 2010, 2011). The Hfq protein, which is encoded from one of the most highly upregulated genes in the *sif* mutant strain, could inhibit the translation of A118 phage genes and therefore promote active lysogeny. Noteworthy, the Hfq protein was originally identified as HF-I - a host factor for phage Qbeta RNA replication in *E. coli* (Franze de Fernandez et al., 1968), which further supports a putative link between Hfq and A118 phage of *L. monocytogenes*. An intriguing possibility is also the potential involvement of the Zea protein in the post-transcriptional control of A118 phage gene expression. The Zea protein has been described very recently as the protein specifically binding transcripts of A118 phage genes of *L. monocytogenes* EGD-e (Pagliuso et al., 2019). Unfortunately, it is not known whether Zea binding to the phage RNAs influences their translation, however the observed decrease in *zea* expression in the *sif* mutant strain may suggest that it is involved in the control of active lysogeny of A118 phage in the EGD-e strain. Clearly, our speculations on the potential involvement of Hfq and/or Zea proteins in post-transcriptional regulation of A118 phage genes expression require further research.

Our study also revealed that *L. monocytogenes* responds to *lmo0946* inactivation by upregulation of the monocin genetic locus. As the monocin element in *L. monocytogenes* is reported to be activated under UV irradiation (Argov et al., 2017), we assume that high level of expression of this MGE in the *sif* imutant strain, similarly to phage A118, results from activation of the SOS response. It is worth noting that the monocin locus resembles a phage tail module of A118-like phages but it does not contain the DNA region responsible for DNA replication and recombination (Lee et al., 2016). These observations might explain why this MGE element, unlike phage A118, is not mobilized. Inactivation of the *sif* gene also results in replication of ICELm1 and the presence of this element in circular form in *L. monocytogenes* EGD-e. Generally, integrative and conjugative elements (ICEs), also known as conjugative transposons, are normally integrated into the bacterial chromosome. They can excise, transfer by conjugation and integrate into the chromosome of the recipient (Whittle et al., 2002). Notably, this is the first report on mobilization of the ICE in *L. monocytogenes*. In case of the integrative and conjugative element ICE*Bs1* of *Bacillus subtilis*, capable of transferring to *Bacillus* and *Listeria* species, its excision and transfer is activated by induction of the SOS response (Auchtung et al., 2005). Therefore, we assume that the replication and presence of ICELm1 in circular form in the *sif* mutant strain may be associated with activation of the SOS response which results from *sif* inactivation. The replication and presence of ICELm1 in circular form in the *sif* mutant strain is correlated with increased resistance to cadmium, which confirms the mobilization of this MGE. Curiously, upon complementation ICELm1 is still detected in a circular form and the presence of this MGE in the complemented strain correlates well with high level of expression of *lmo1097* coding for an integrase of ICELm1 and high level of cadmium resistance. Thus, ICELm1 mobilization and associated phenotype is not reversed upon complementation. Moreover, the complementation also fails to restore the expression of several other genes, namely *hfq*, *zea* and *rsbV*, to the level observed in the wild-type strain. At the current stage of research, the reason for the lack of reversion of the expression level of these genes and the mobilization of ICELM1 upon complementation remains unclear. However, we suppose that the lack of complementation may be due to a lower level of Sif protein in the complemented strain as compared to the wild-type strain. In the complemented strain, the *sif* gene is expressed *in trans* from the native promoter of *lmo0945.* As a result, Sif is not translated from the larger co-transcripts corresponding to *sif* mRNA arising in *L. monocytogenes* EGD-e, which in turn leads to an overall lower level of Sif protein in the complemented strain than in the wild-type strain.

Activation of the SOS response and the mobilization of MGEs, is accompanied by a decrease in energy production and conversion as well as transport and metabolism of different cell compounds in the *sif* mutant strain. These changes indicate general metabolic slowdown in response to *sif* inactivation. Our analysis also revealed that *L. monocytogenes* responds to *sif* inactivation by changed expression of multiple genes known to be under control of important regulators involved in stress response and virulence of *L. monocytogenes,* including CodY, σB and PrfA. The largest overlap of genes with highly changed expression in the *sif* mutant is observed for the CodY regulon. The transcriptional regulator CodY plays an important role in carbon and nitrogen assimilation and methabolism. The role of CodY relies on monitoring the overall energetic state of the cell by sensing the levels of GTP, and the nutritional state of the cell by sensing the levels of branched chain amino acid (BCAAs) (Bennett et al., 2007). In the current study, no change in the level of expression of the BCAA biosynthesis operon (*ilv-leu*) was observed (Supplementary Table S2). This suggests that putative modulation of CodY activity in the *sif* mutant strain does not result from BCAAs insufficiency and could be rather GTP-dependent. This assumption is in line with the observed general downregulation of genes involved in energy production and conversion in the *sif* mutant strain. However, it should be noted that only some of the genes overlapping with the CodY regulon are involved in transport and metabolism, while most of them correspond to A118 prophage genes. Therefore, the involvement of CodY in the downregulation of energy production and conversion in the *sif* mutant strain is uncertain. Similarly, the engagement of CodY in the regulation of A118 prophage genes in the *sif* mutant is ambiguous, as the details of CodY-dependent regulation of A118 prohage genes are missing (Bennett et al., 2007), and upregulation of these genes is triggered by activation of the SOS response (Pasechnek et al., 2020).

Among the highly down-regulated genes in the *sif* mutant strain, we found genes positively regulated by the alternative sigma factor σB. The σB factor controls the general stress response in *L. monocytogenes* (Dorey et al., 2019) and plays an indespensible role in the gastrointestinal phase of infection (Toledo-Arana et al., 2009). One of the stresses experienced by *L. monocytogenes* in the gastrointestinal tract is bile stress, and σB-dependent expression of *bsh* encoding the bile salt hydrolase enables bile tolerance (Begley et al., 2005). The downregulation of *bsh* in the *sif* mutant strain suggests that inactivation of *sif* leads to an impairment of stress tolerance during the gastrointestinal phase of infection. Further analysis revealed decreased level of expression of *rsbV* which is an important factor in the signaling pathway controlling σB activity. In stress-free conditions, σB is sequestered by the anti-sigma factor RsbW that prevents interaction of σB with core RNA polymerase, and therefore the σB regulon is not transcribed (Ferreira et al., 2001). When stress signals are sensed, the anti-anti-sigma factor RsbV undergoes dephosphorylation and interacts directly with RsbW. The RsbV-RsbW interaction results in release of σB from RsbW-σB complexes. Subsequently, σB binds to the core RNA polymerase and directs transcription of the σB regulon (Hecker et al., 2007). Therefore, RsbV is required to activate σB in response to stress signals in *L. monocytogenes* (Chaturongakul & Boor, 2004, 2006). It seems likely that the reduced level of *rsbV* expression in the *sif* mutant strain may result in decreased liberation of σB from the RsbW-σB complexes, and thus in inhibition of transcription of the σB regulon despite of stress signals sensed. While the impairment of the σB signaling pathway does not explain the downregulation of numerous genes which are under positive control of σB, it clearly indicates that the σB protective response is prevented in *sif* mutant strain. Interestingly, while *rsbV* is the first gene of the four-gene operon which includes besides *rsbV* also *rsbW*, *sigB* and *rsbX*, we observed that expression of *rsbV* but not *sigB* is downregulated. This result suggests that the expression of this operon is regulated post-transcriptionally in the *sif* mutant strain.

Finally, we found that expression of a large portion of genes regulated by the master virulence regulator PrfA is downregulated in the *sif* mutant strain despite increased expression of *prfA* itself. It should be noted that σB-dependent promoters preceede multiple PrfA-dependent genes which are downregulated in the *sif* mutant strain (Supplementary Table S7, Milohanic et al., 2003). This suggests that downregulation of PrfA-dependent genes is related to the downregulation of σB-dependent genes described above for the *sif* mutant strain. However, among the downregulated genes are also those belonging to the core set of genes preceded by a PrfA box (e.g., *hly* encoding listeriolysin O); these genes are most likely expressed in a σA-dependent manner (Milohanic et al., 2003). Furthermore, among highly downregulated genes is *lmo1634*, encoding the LAP protein with a role in crossing the intestinal epithelial barrier (Drolia et al., 2018) as well as *arpJ* and *pplA* coding for factors important at the intracellular stage of infection (Reniere et al., 2016). Interestingly, the virulence factors mentioned above do not belong to the PrfA- or σB regulons. These findings indicate that inactivation of *sif* impairs virulence of *L. monocytogenes* at different stages of infection. This conclusion is supported by the diminished virulence of the *sif* mutant observed in our *in vitro* and *in vivo* infection studies. While the details of the complex regulatory circuits in the *sif* mutant strain remain to be elucidated, it is clear that inactivation of *sif* leads to general silencing of the virulence program in *L. monocytogenes*.

To summarize, we found that *fri* is transcribed together with four downstream genes, namely *lmo0944*, *lmo0945, lmo0946* and *lhrC5*. Functional analysis of genes from the ferritin operon revealed that inactivation of *lmo0946*, coding for a protein with unknown function, results in severe impairment of *L. monocytogenes* growth in virualy all conditions tested. Inactivation of *lmo0946* results in global changes of gene expression with clearly induced expression of SOS response genes, which prompted us to rename the *lmo0946* gene to *sif* (<u>S</u>OS interfering factor). Activation of the SOS response in the *sif* mutant strain is accompanied by mobilization of A118 prophage and ICELm-1 mobile genetic elements (MGEs). In parallel to activation of the SOS response and mobilization of MGEs, the absence of Sif results in downregulation of stress response genes from the σ^B^ regulon and multiple virulence genes, modulation of expression of genes from the CodY regulon, and highly upregulated expression of *hfq* coding for an RNA chaperone. While the function of Sif could not be disclosed in the current study, the results clearly demonstrate the importance of this newly discovered protein in stress adaptation and virulence of *L. monocytogenes*. Future studies should aim at disclosing the function of Sif as well as defining the molecular mechanisms responsible for the transcriptional changes resulting from *sif* inactivation, especially those related to silencing of the general stress response and virulence program in *L. monocytogenes*.

## MATERIALS AND METHODS

### Bacterial Strains and Growth Conditions

The wild-type strain *L. monocytogenes* EGD-e, its derivative mutant strains in ferritin operon genes (described in detail below) and strain with complemented mutation of *lmo0946* were used in this study. *L. monocytogenes* was routinely grown in brain heart infusion broth (BHI, Oxoid) at 37 ◦C with aeration. When appropriate, cultures were supplemented with chloramphenicol (7.5 μg/ml), erythromycin (5 µg/ml) or X-gal (50 mg/ml).

For growth experiments, overnight cultures grown at 37°C with aeration in BHI medium were diluted 1:1000 in fresh BHI broth in titration plates. Depending on the experiment, BHI medium was suplemented with 5 % ethanol, 0.09 µg/ml penicillin G, acidified with HCl to pH 5 or alkalized with NaOH to pH 9. The cultures were grown at 37°C with aeration and bacterial growth was monitored by measuring the optical density at 600 nm (OD_600_). For thermal stress, the cultures were grown at 37°C to an OD_600_ of 0.03 and at this point the incubation temperature was shifted to 43°C. *Escherichia coli* strain Top10 and BL21(DE3) (Invitrogen) were used in cloning experiments and purification of Lmo0946-His_6_, respectively. *E. coli* strains were grown on Luria-Bertani medium. When required, the LB medium was supplemented with ampicillin (100 μg/ml), chloramphenicol (25 μg/ml) or kanamycin (30 µg/ml).

### Construction of ferritin operon mutant strains and complementation of *lmo0946* mutation

For the construction of in-frame mutants with deletions of *lmo0944*, *lmo0945*, *lhrC1-4*, *lhrC6* (*rli22*) and *lhrC7* (*rli33.1*) *L. monocytogenes* EGD-e chromosomal DNA was used as the template for the PCR amplification of DNA fragments representing either the 5’ end and upstream sequences or the 3’ end and downstream sequences of the respective genes. The Splicing of the amplified 5’ and 3’ fragments were performed by overlap extension PCR and the obtained products were subsequently cloned into the temperature-sensitive shuttle vector pMAD (Arnaud et al., 2004). For the construction of nonsense mutants of *fri* and *lmo0946* and mutant with disabled -10 promoter region of *lhrC5 L. monocytogenes* EGD-e chromosomal DNA was used as the template for the PCR amplification of DNA fragments covering intended mutagenesis site of the respective genes. The obtained PCR products were cloned into the temperature-sensitive shuttle vector pMAD and obtained derivatives of pMAD vector were used as the tempaltes for site-directed mutagenesis (SDM). In case of *fri*, the introduced nucleotide substitutions changed the TTA codon at position 58-60 bp of the open reading frame to the STOP codon TAG. In case of *lmo0946*, the introduced nucleotide substitutions changed the GAA codon at position 70-72 bp of the open reading frame of *lmo0946* to the STOP codon TAG. In case of *lhrC5*, the introduced nucleotide substitutions changed the -10 promoter consensus sequence TATTAT to CAGTGC. Primers used for constructing the in-frame deletions as well as site-directed mutagenesis are listed in Supplementary Table S9. The obtained pMAD derivatives were used for genes replacement which was performed via double-crossover homologous recombination as described previously (Arnaud et al., 2004). Erythromycin-sensitive clones were screened for the presence of the mutations by PCR. Accuracy of the desired DNA modifications in the obtained mutant strains was confirmed by the sequencing of obtained PCR products.

For the construction of the strain with complemented *lmo0946* mutation *L. monocytogenes*Δ*lmo0945* chromosomal DNA was used as the template for the PCR amplification of DNA fragment covering ORFs of *lmo0945* and *lmo0946.* The obtained PCR product was cloned into the pPL2 vector and subsequently introduced into *lmo0946\** mutant strain as described previously (Lauer et al., 2002).

### Biofilm Formation analysis

Biofilm formation was carried out essentially as described previously (Djordjevic et al., 2002). Briefly, 12-well polystyrene plates containing 1 ml of BHI broth were inoculated with the overnight culture of each strain to OD_600_ = 0.01. The plates were then incubated at 37°C under static condition. At different time points (14, 24, and 72 h), the wells were emptied and washed with sterile distilled water. After fixation with methanol, the wells were air dried and stained with crystal violet solution for 5 min. After washing, wells were air dried and the bound dye was solubilized with an solution of acetic acid. Following 10-fold dilution in destiled wated, the absorbance of the solubilized dye was measured at 570 nm.

### Susceptibility tests

The susceptibility to antibiotics and cadmium as well as the tolerance to penicillin G were examined essentially as described previously (Krawczyk-Balska et al., 2012). Briefly, The susceptibility to antibiotics and cadmium were examined using a microdilution test. 96-well microdilution plates containing 100 μl of two-fold dilutions of antibiotics or cadmium chloride (Merck) in BHI broth were innoculated with 10^5^ CFU/ml of overnight culture of each strain. The plates were then incubated at 37°C. The MIC endpoints were read after 18–22 h of incubation for antibiotics susceptibility testing and after 48 h of incubation for cadmium susceptibility testing. The MIC was determined as the lowest tested compound concentration that resulted in the absence of apparent growth of the bacteria.

The tolerance to penicillin G was examined by innoculation of BHI broth supplemented with 32 μg/ml penicillin G with 5 × 10^7^ CFU/ml of mid-exponential phase cultures. The cultures were subsequently incubated with aeration at 37°C and at indicated points of time viable cell counts were performed by standard plate counting.

### In vitro infection experiment

Infection experiments with *L. monocytogenes* strains were performed using P388.D1 murine macrophage (ECACC Collection) essentially as described previously (Chatterjee et al., 2006). Briefly, macrophages were grown in 12-well plates until reached at least 80% confluence. Bacteria were added at MOI (multiplicity of infection) of 10. After 45 minutes remaining extracellular bacteria were removed and macrophages were washed twice with 1 x phosphate buffered saline (PBS), and subsequently incubated with fresh medium suplemented with gentamycin (20 µg/ml). After 1 and 4 hours post infection macrophages were washed three times with 1x PBS, lysed in 0.1% Triton X-100 and viable cell counts were performed by standard plate counting.

### Murine infection model

*In vivo* infection experiments were performed essentially as described previously (Cabanes et al., 2008). Briefly, 2*10^3^ bacteria in 200 μl of 1xPBS were administered intravenously to 6-8 old BALB/c female mice. After 72h post infection bacterial growth in mice spleens and livers was determined by plating 10-fold serial dilutions of organ homogenates on BHI.

### Ethics Statement

All animal procedures complied with the Act on the Protection of Animals used for scientific or educational purposes dated on 15th of January 2015 (D20150266L), which implements the Directive of the European Parliament and the Council (2010/63/EU). The animal procedures were in agreement with the guidelines of the Institutional Animal Care and Use Committee (IACUC) and the whole study was approved by 2nd Local IACUC in Kraków (Institute of Pharmacology Polish Academy of Sciences, Permit Number: 48/2017).

### Total RNA Isolation and Purification

For northern blot analysis in stationary phase, *L. monocytogenes* EGD-e was innoculated from single colony into BHI broth and cultured for 18h. For northern blot analysis in exponential phase, *L. monocytogenes* EGD-e was grown to OD_600_ ∼ 0.35, split, and half stressed with 0.09 µg/ml penicyllin G for 60 min. 10 ml samples from stationary phase cultures and 20 ml from exponential phase cultures were immedietely cooled down in liquid nitrogen and centrifuged at 12 000 x g for 3 min at 4◦C. The cell pellets were frozen in liquid nitrogen and stored at - 80 °C until further processing. The cells were disrupted by the FastPrep instrument and total RNA was extracted using 1 ml of TRItidy G™ Reagent (Applichem) as described previously (Nielsen et al., 2010). The concentration and purity of RNA were determined with a NanoDrop ND-1000 spectrophotometer and integrity of RNA was confirmed by agarose gel electrophoresis.

For transcriptomic and RT-qPCR analysis, *L. monocytogenes* cultures were grown to OD_600_ ∼ 0.35. 5 ml samples of each culture were stabilized with RNA protect Cell Reagent (Qiagen) and subsequently collected by centrifugation at 5 000 x g for 10 min at 4◦C. The cell pellets were frozen immediately in liquid nitrogen and stored at - 80 °C until further processing. The pellets were suspended in 1 mL of TRItidy G™ Reagent (Applichem) and cells were disrupted by the FastPrep instrument. Total RNA was isolated with Direct-zol RNA MiniPrep (Zymo Research) according to the manufacturer’s instructions. Contaminating DNA was removed from the samples using the TURBO DNA-free™ Kit (Invitrogen) following the manufacturer‘s instructions, except that DNAse treatment was performed in presence of SUPERase•In™ RNase Inhibitor (Invitrogen). The concentration and purity of the RNA preparations were then estimated by measuring the A260 and A280 with a NanoDrop ND-1000 spectrophotometer. The absence of DNA from RNA preparations was verified by the failure to amplify a *rpoB* gene fragment in a 30-cycle PCR using 100 ng of each RNA isolation as the template. A 2100 Bioanalyzer (Agilent Technologies) was used to measure the RNA quality. The prepared RNA was stored at −80°C before further analysis.

### rRNA Depletion, Library Preparation, RNA Sequencing and Data Analysis

Ribosomal RNA depletion, library preparation and sequencing were performed by Eurofins Genomics AT GmbH (Vienna, Austria). To enrich mRNA and remove ribosomal RNA (rRNA) from total RNA, total RNA was treated with Bacterial rRNA Depletion Kit (NEB). Preparation of cDNA fragment libraries was performed using the NEBNext^®^ Ultra^TM^ II Directional RNA Library Prep Kit for Illumina^®^ (Illumina). Sequencing was carried out on the Illumina NovaSeq platform (10 M paired-end reads, 2×150 bp per read). The obtained raw data were further analyzed by DDG Bioinformatics, University of Warsaw. Briefly, sequenced reads were quality-checked using FastQC, and Illumina adapter sequences and low-quality base pairs were removed using fastp (Chen et al., 2018). Reads were mapped against the complete sequenced genome of the *L. monocytogenes* reference strain EGD-e (ENSEMBL ASM19603v1) and 154 genes of ncRNA from Listeriomics database, using Bowtie 2 with standard parameters and sensitive-local (Langmead & Salzberg, 2012). Differential expression (DE) analyses were performed using DESeq2 (Love et al., 2014). DE was reported as log_2_fold changes and p-values were adjusted by the DESeq2 default Benjamini–Hochberg (BH) adjustment method. The reading counts obtained from the Salmon program (Patro et al., 2017) were submitted to the DEseq2 tool using the tximport (Soneson et al., 2015) library. The results of the differential gene expression analysis were additionally scaled using the apeglm library (Zitovsky & Love, 2019), the aim of which was to reduce the influence of high variability of low-expressed genes on the results of the analysis. The fold changes were calculated by comparing expression levels of *lmo0946* mutant strain vs. wild-type strain. Genes with changed expression and an adjusted p-value <0.01 were considered as DE in the interpretation applied in this study. The transcriptomic data has been deposited on the Array Express database and is available via the Experiment ArrayExpress accession E-MTAB-10809.

### Reverse Transcriptase-Quantitative Polymerase Chain Reaction (RT-qPCR)

100 ng of total RNA was used for synthesis of first strand cDNA. Reverse transcription was performed with RevertAid First Strand cDNA Synthesis Kit and random hexamers used as primers (Thermo Fisher Scientific) according to the manufacturer’s protocol. Quantitative PCR reactions were performed using the LightCycler^®^ 480 SYBR Green I Master (Roche) and specific primer sets for the gene of interest (Supplementary Table S9). The samples were run on LightCycler^®^96 thermocycler (Roche) with an initial step at 95◦C for 10 min, 45 cycles of 10 s at 95◦C, 10 s at 60◦C and 15 s at 72◦C. Data was analyzed using LightCycler® 96 SW 1.1 Software. The relative amount of target cDNA was normalized using the *rpoB* gene. The experiment was carried out in three biological replicates, each in technical duplicates.

### Co-transcription and Northern Blotting Analysis

For co-transcription analysis, 100 ng of total RNA was used for synthesis of first strand cDNA. Reverse transcription was performed with RevertAid First Strand cDNA Synthesis Kit (Thermo Fisher Scientific) and primer specific for the *lhrC5* gene. The obtained cDNA was then used as the template for PCR performed with primers specific for internal fragments of the *fri* and *lmo0944* genes. Primers used for the co-transcription analysis are listed in Supplementary Table S9.

For northern blotting, 20 µg of total RNA in loading buffer containing 50% formamide and 20% formaldehyde was separated on a formaldehyde agarose gel and subsequently transferred to a Zeta probe nylon membrane (Bio-Rad) by capillarity blotting. For detection of RNA, the membranes were preincubated for 1h in PerfectHyb hybridization buffer (Sigma-Aldrich) and then hybridized overnight with a specific ^32^P-labelled DNA double-stranded probes. Probes were generated using [α-32P]dATP and Megaprime DNA labeling system (Amersham Biosciences), according to the manufacturer’s protocol. Primers used for preparing DNA double-stranded probes are listed in Supplementary Table S9. RNA bands were visualized by phosphor imaging using a Typhoon scanner and analyzed with ImageQuant™ TL software (GE Healthcare).

### MGE mobilization analysis

15 ml of exponential phase cultures (OD_600_∼0.35) were centrifuged at 5 000 x g for 10 min. The cell pellets were suspened in buffer containing 0.2 mg/ml lysozyme and 20 % saccharose and incubated at at 37°C for 5 min. Total DNA was subsequently isolated with GeneJetGenomic DNA Purification Kit (ThermoScientific) according to the manufacturer’s instructions. The concentration of DNA was determined with a NanoDrop ND-1000 spectrophotometer. PCR reactions were performed using 0.1 ng of DNA and specific primer sets (Supplementary Table S9).

### Purification of Sif protein and Electrophoretic mobility shift assay (EMSA)

The His-tag system (Novagen) was used for Sif purification. The fragment representing the entire *lmo0946* coding sequence was PCR-amplified on the template of *L. monocytogenes* EGD-e chromosomal DNA using primers listed in Supplementary Table S9. PCR product was cloned into vector pET28a (Novagen) and the obtained construct was used to transform *E. coli* BL21(DE3). The N-terminal His-tagged Sif protein (Lmo0946-His_6_) was expressed and purified using Ni-NTA resin (Qiagen) according to the manufacturer’s recommendations. Briefly, the cells were suspended in 20 mM Tris-HCl (pH 8.0) buffer containing 500 mM NaCl, 1 mM PMSF and 5 mM imidazol and disrupted by sonication. After centrifuging the cell lysate to remove unbroken cells, the supernatant was passed through a Ni-NTA agarose column. The column was washed with 20 mM Tris-HCl (pH 8.0) buffer containing 500 mM NaCl, 10% glicerol, 5 mM imidazol, and subsequently the Lmo0946-His_6_ protein was eluted using a gradient of imidazole buffer. Finally, the protein was dialyzed against 20 mM Tris-HCl (pH 8.0), 150 mM NaCl and 1% glicerol. The concentration of the purified Sif was determined using the Pierce™ BCA Protein Assay Kit (ThermoScientific).

For EMSA analysis, DNA fragments comprising the regulatory regions upstream of genes of interest were PCR-amplified using primers listed in Supplementary Table S9. The amplicons were purified using a GeneJET Gel Extraction & DNA Cleanup Micro Kit (ThermoScientific) and the concentration of DNA was determined with a NanoDrop ND-1000 spectrophotometer. EMSA reactions contained appropriate DNA fragments of the studied promoters (0.1 pmol) and Lmo0946-His_6_ in concentration ranging from 0 to 40 μM. The purified protein was used in EMSAs with and without *in vitro* phosphorylation. A 202-bp fragment of *L. monocytogenes* 16S rDNA generated by PCR was included as a non-specific competitor in binding reactions. Sif binding to the DNA targets was performed at 37 °C for 30 min and subsequently samples were separated on 4.2% native polyacrylamide gels. After electrophoresis the gels were stained with ethidium bromide and DNA bands were visualized using a AI600 imager (GE Healthcare).

### Competing interests

The authors declare no competing interests. The funders had no role in the design of the study; in the collection, analyses, or interpretation of data; in the writing of the manuscript, or in the decision to publish the results.

### Authors’ contributions

Conceptualization A.K-B, B.H.K; data curation: A.K-B; formal analysis: A.K-B, M.Ł; funding acquisition: A.K.B; investigation: M.Ł, E.P, K.G, P.G, K.J, K.Ś, M.M, K.K, A.K-B; methodology: M.Ł, E.P, K.G, K.J, E.M.S.L, K.P, B.H.K, A.K-B; project administration: A.K-B; resources: K.P, B.H.K, A.K-B; supervision: A.K-B; validation: M.Ł, P.G, K.J; visualisation: K.J, A.K-B; writing—original draft preparation M.Ł, B.H.K, A.K.B; writing—review and editing A.K-B, B.H.K

## Acknowledgments

The study was supported by the National Science Centre (Poland) through a grant to AKB (grant no. 2016/21/B/NZ6/00963) and partially supported by EMBO Short-term Fellowship (no. ASTF 137) to AKB. The authors are grateful to Krzysztof Pyrć from Malopolska Centre of Biotechnology, Jagiellonian University for the organizational care over animal experiments and Inga Drebot from Malopolska Centre of Biotechnology for expert technical assistance in *in vivo* infection experiments. The authors thank also Marta Zapotoczna for help with biofilm analysis, Mikołaj Dziurzyński from DDG Bioinformatics for great expert assistance in transcriptomics data analysis, and Dariusz Bartosik from Department of Bacterial Genetics, University of Warsaw for a constructive discussion on MGEs.

## Notes

### Competing Interest Statement

The authors have declared no competing interest.

