## Supplementary figures and images for "Inactivation of *lmo0946* (*sif*) induces the SOS response and MGEs mobilization and silences the general stress response and virulence program in *Listeria monocytogenes*"

### Supplementary Figure S1

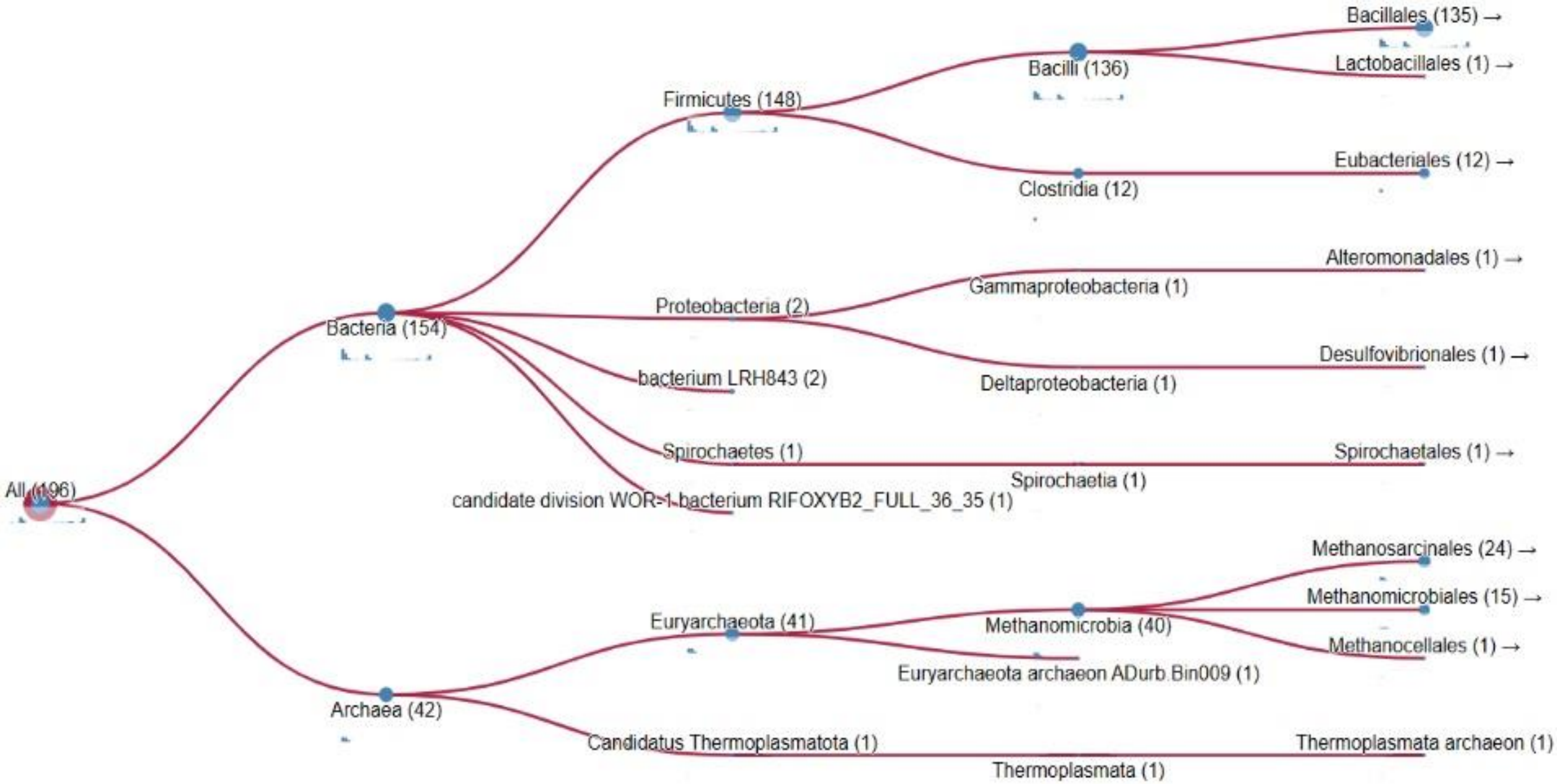


**Figure S1. PHAMMER analysis of taxonomic distribution of Lmo0946 homologs in UniProtKB database.**
