## Supplementary Figure S2 for "Inactivation of *lmo0946* (*sif*) induces the SOS response and MGEs mobilization and silences the general stress response and virulence program in *Listeria monocytogenes*"

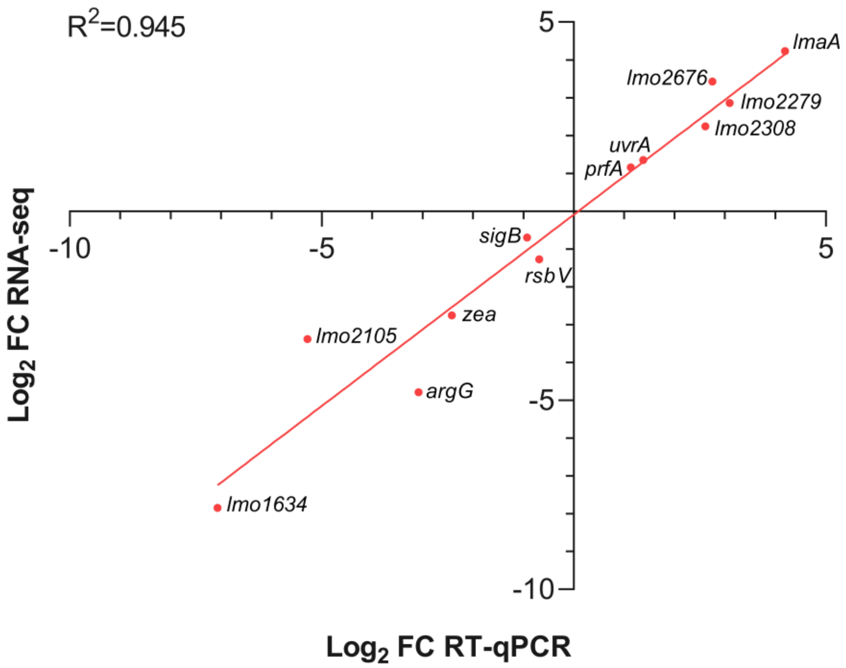


**Figure S2. Validation of the RNA sequencing data by RT-qPCR analysis. Comparison of the log2 of the fold changes (FC) of 12 genes obtained in the RNA-seq and in the RT-qPCR.** The data represent mean values from 3 biological replicates of experiments.
