## Supplementary Figure S3 for "Inactivation of *lmo0946* (*sif*) induces the SOS response and MGEs mobilization and silences the general stress response and virulence program in *Listeria monocytogenes*"

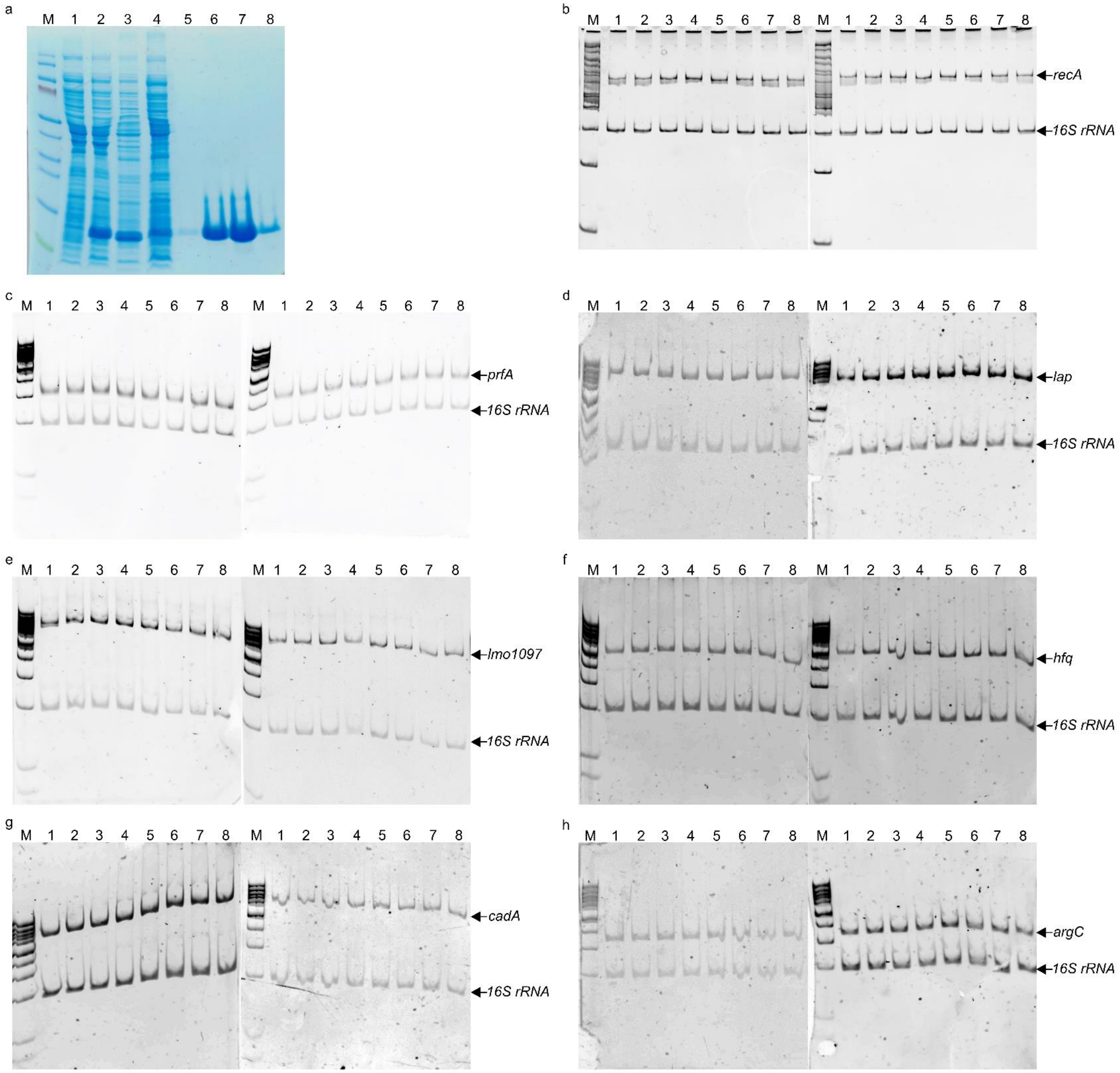


**Figure S3. EMSA analyses of the interaction between selected genes regulatory regions and Lmo0946-His$_6$ protein.** (**a**) Coomassie blue SDS-Page showing steps of His-tag/nickel column purification of Lmo0946-His$_6$ protein. M – Page Ruler Prestained Protein Ladder; 2 – total cell extract before induction of expression with IPTG; 3 – total cell extract after induction of expression with IPTG; 4 – 4 µl of cell pellet from 2 L bacterial culture; 5 – first fraction after washing of column with 10 mM imidazole buffer; 6 – first fraction after washing of column with 20 mM imidazole buffer; 7 – 16 µl from 18 ml elution fraction of column with 40 mM imidazole buffer; 8 – 16 µl from 18 ml elution fraction of column with 40 mM imidazole buffer; 9 – first fraction after washing of column with 1000 mM imidazole buffer. (**b-h**) EMSA analyses of interaction between phosphorylated (left panels) and unphosphorylated (right panels) Lmo0946-His$_6$ protein and regulatory regions of: b – *recA* (482 bp); c – *prfA* (535 bp); d – *lap* (561 bp); e – *lmo1097* (641 bp); f – *hfq* (455 bp); g – *cadA* (493 bp); h – *argC* (276 bp). 203 bp fragment of *16S rRNA* gene was included in all reaction mixtures and served as negative control. In all cases, the DNA was incubated without protein (lane 1) or with 0.625 (lane 2), 1.25 (lane 3), 2.5 (lane 4), 5 (lane 5), 10 (lane 6), 20 (lane 7), 40 µM (lane 8) Lmo0946-$His_6$. M – Low Range DNA Marker.
