## Supplementary Table S1 for "Inactivation of *lmo0946* (*sif*) induces the SOS response and MGEs mobilization and silences the general stress response and virulence program in *Listeria monocytogenes*"

Table S1. Generation time of *L.monocytogenes* strains in different conditions of growth on the basis of exponential phase of growth analysis.

| | **Generation time [minutes]** | | | | |
|---|---|---|---|---|---|
| Conditions of growth / Strain | 37 °C, BHI (stress-free) | Ethanol 5 % | Penicillin G 0.09 μg/ml | pH 5 | pH 9 |
| EGD-e | 45 | 94 | 54 | 74 | 65 |
| *lmo0946** | 50 | 105 | 62 | 91 | 72 |
| *lmo0946*- lmo0946* | 45 | 89 | 54 | 77 | 61 |
