## Supplementary Table S4 for "Inactivation of *lmo0946* (*sif*) induces the SOS response and MGEs mobilization and silences the general stress response and virulence program in *Listeria monocytogenes*"

**Table S4. Genes ≥ 4.0-fold up-regulated and ≤ - 4.0-fold down-regulated (with Padj <0.01) in *L. monocytogenes* EGD-e *lmo0946** compared to *L. monocytogenes* EGD-e wild-type strain in cultures from the exponential phase of growth in BHI in 37°C.**

| Gene name | Gene symbol | $log_2$ Fold Change | Regulation[a] | Product[b] | COG[b] |
|---|---|---|---|---|---|
| *lmo1097* | | 13.71 | | Integrase, superantigen-encoding pathogenicity islands SaPI | Replication, recombination and repair |
| *lmo1295* | | 12.65 | **SigB** [1] ↑ | RNA-binding protein Hfq | Transcription; Translation |
| *lmo1100* | *cadA* | 12.27 | | Cadmium resistance protein | Inorganic ion transport and metabolism |
| *lmo1101* | *lspB* | 9.12 | | Hypothetical protein | Cell wall/membrane biogenesis; Intracellular trafficking and secretion |
| *lmo1102* | *cadC* | 7.85 | | Cadmium efflux system accessory protein | Transcription |
| *lmo1105* | | 6.36 | | Membrane protein, putative | Not in COGs |
| *lmo1112* | | 5.29 | | Hypothetical protein | Cell cycle control, mitosis and meiosis |
| *lmo1106* | | 5.18 | | Hypothetical protein | Not in COGs |
| *lmo1115* | | 4.96 | | Similar to fibrinogen-binding protein | Cell wall/membrane biogenesis |
| *lmo2325* | | 4.81 | | Hypothetical protein | Not in COGs |
| *lmo0119* | | 4.61 | | Hypothetical protein | Not in COGs |
| *lmo0117* | *lmaB* | 4.61 | | Antigen B | Not in COGs |
| *lmo2294* | | 4.53 | **CodY** [7] ↑ | Protein gp9 | Signal transduction mechanisms |
| *lmo0118* | *lmaA* | 4.52 | | Antigen A | Function unknown |
| *lmo0120* | | 4.50 | | Hypothetical protein | Not in COGs |
| *lmo2321* | | 4.35 | | Protein gp45 [Bacteriophage A118] | Not in COGs |
| *lmo2289* | | 4.35 | | Protein gp14 | Not in COGs |
| *lmo0126* | | 4.32 | | Hypothetical protein | Not in COGs |
| *lmo0121* | | 4.28 | | Phage tail length tape-measure protein | Function unknown |
| *lmo0125* | | 4.26 | | Hypothetical protein | Not in COGs |
| *lmo0127* | | 4.24 | | Hypothetical protein | Not in COGs |
| *lmo0122* | | 4.23 | | Phage tail fiber | Not in COGs |
| *lmo2292* | | 4.19 | **CodY** [7] ↑ | Protein gp11 | Not in COGs |
| *lmo0123* | | 4.15 | | Putative tail or base plate protein gp18 [Bacteriophage A118] | Not in COGs |

| | | | | | |
|---|---|---|---|---|---|
| *lmo2324* | | 4.12 | **CodY** [7] ↑ | Phage antirepressor protein / Antirepressor [Bacteriophage A118] | Transcription; Function unknown |
| *lmo0129* | | 4.12 | | N-acetylmuramoyl-L-alanine amidase | Cell wall/membrane biogenesis |
| *lmo0128* | | 3.99 | | Similar to phage-related protein | General function prediction only |
| *lmo0124* | | 3.96 | | Hypothetical protein | Not in COGs |
| *lmo2315* | | 3.90 | | Protein gp51 [Bacteriophage A118] | Not in COGs |
| *lmo2316* | | 3.88 | | Methyltransferase | Replication, recombination and repair |
| *lmo2326* | | 3.82 | | Protein gp41 [Bacteriophage A118] | Not in COGs |
| *lmo2298* | | 3.81 | **CodY** [7] ↑ | Protein gp4 | Not in COGs |
| *lmo2313* | | 3.75 | **CodY** [7] ↑ | Hypothetical protein, Lmo2313 homolog [Bacteriophage A118] | Not in COGs |
| *lmo0115* | *lmaD* | 3.74 | | Listeria protein LmaD, associated with virulence | Not in COGs |
| *lmo2288* | | 3.74 | **CodY** [7] ↑ | Protein gp15 | Not in COGs |
| *lmo2676* | *umuC* | 3.70 | **LexA/RecA** [3] ↑<br>**CodY** [7] ↓ | Hypothetical protein | Replication, recombination and repair |
| *lmo2675* | *umuD* | 3.70 | **LexA/RecA** [3] ↑<br>**CodY** [7] ↓ | Hypothetical protein | Not in COGs |
| *lmo2291* | | 3.68 | **CodY** [7] ↑ | Major tail shaft protein | Not in COGs |
| *lmo0116* | *lmaC* | 3.66 | | LmaC, associated with virulence in Listeria | Not in COGs |
| *lmo2322* | | 3.64 | | gp44 | Not in COGs |
| *lmo2296* | | 3.62 | **CodY** [7] ↑ | Phage capsid protein | Not in COGs |
| *lmo2327* | | 3.59 | | Hypothetical protein | Not in COGs |
| *lmo2282* | | 3.59 | **CodY** [7] ↑ | protein gp21 | Not in COGs |
| *lmo2323* | | 3.58 | | Protein gp43 [Bacteriophage A118] | Not in COGs |
| *lmo2318* | | 3.52 | **CodY** [7] ↑ | Putative recombination protein / Single-stranded DNA-binding protein | Not in COGs |
| *lmo2271* | | 3.51 | **LexA/RecA** [3] ↑ | Hypothetical protein | Not in COGs |
| *lmo2285* | | 3.51 | **CodY** [7] ↑ | Protein gp18 | Translation |
| *rli38* | | 3.51 | | | |
| *lmo2319* | | 3.50 | | Hypothetical protein | Not in COGs |
| *lmo2299* | | 3.46 | **CodY** [7] ↑ | Putative portal protein | Not in COGs |
| *lmo2293* | | 3.39 | **CodY** [7] ↑ | Protein gp10 | Not in COGs |

| | | | | | |
|---|---|---|---|---|---|
| *lmo2317* | | 3.35 | **CodY** [7] ↑ | Protein gp49, replication initiation [Bacteriophage A118] | Replication, recombination and repair |
| *lmo2300* | | 3.33 | **CodY** [7] ↑ | Putative terminase large subunit from bacteriophage A118 | General function prediction only |
| *lmo2295* | | 3.30 | **CodY** [7] ↑ | Protein gp8 | Not in COGs |
| *lmo2297* | | 3.29 | **CodY** [7] ↑ | Putative scaffolding protein | Cell motility; Signal transduction mechanisms |
| *lmo2290* | | 3.24 | **CodY** [7] ↑ | Protein gp13 | Cell motility |
| *lmo2287* | | 3.20 | **CodY** [7] ↑ | Putative tape-measure | Not in COGs |
| *lmo0152* | | 3.20 | **VirRS** [8] ↑ | Oligopeptide ABC transporter, periplasmic oligopeptide-binding protein | Amino acid transport and metabolism |
| *lmo2301* | | 3.14 | **CodY** [7] ↑ | Terminase small subunit [Bacteriophage A118] | Replication, recombination and repair |
| *rli140* | | 3.14 | | | |
| *lmo2278* | *lysA* | 3.09 | **CodY** [7] ↑ | L-alanoyl-D-glutamate peptidase | Not in COGs |
| *lmo2284* | | 3.09 | **CodY** [7] ↑ | Protein gp19 | Not in COGs |
| *lmo2303* | | 3.06 | **CodY** [7] ↑ | Protein gp66 | Not in COGs |
| *lmo2320* | | 2.88 | | Hypothetical protein | Not in COGs |
| *lmo2283* | | 2.84 | **CodY** [7] ↑ | Protein gp20 | Not in COGs |
| *lmo2305* | | 2.80 | **CodY** [7] ↑ | Hypothetical protein, Lmo2305 homolog [Bacteriophage A118] | Not in COGs |
| *lmo2312* | | 2.79 | | Conserved hypothetical protein | Function unknown |
| *lmo2279* | | 2.78 | **CodY** [7] ↑ | Holin | Not in COGs |
| *lmo2286* | | 2.75 | **CodY** [7] ↑ | Protein gp17 | Not in COGs |
| *lmo0135* | | 2.65 | **VirRS** [8] ↑ | Oligopeptide ABC transporter, periplasmic oligopeptide-binding protein | Amino acid transport and metabolism |
| *lmo2352* | | 2.63 | | HTH-type transcriptional regulator YtlI, LysR family | Transcription |
| *lmo2828* | | 2.58 | **LexA/RecA** [3] ↑ **CodY** [7] ↓ | Hypothetical protein | Not in COGs |
| *lmo2311* | | 2.46 | | Hypothetical protein | Not in COGs |
| *lmo2280* | | 2.45 | **CodY** [7] ↑ | Protein gp23 | Not in COGs |
| *lmo2308* | | 2.34 | | Single-stranded DNA-binding protein (prophage associated) | Replication, recombination and repair |
| *lmo2306* | | 2.34 | **CodY** [7] ↑ | Hypothetical protein | Not in COGs |
| *lmo1975* | *dinB* | 2.29 | **LexA/RecA** [3] ↑ | DNA polymerase IV | Replication, recombination and repair |
| *lmo2744* | | 2.23 | **CodY** [7] ↓ | Cyclic nucleotide-binding protein | Signal transduction mechanisms |

| | | | | | |
|---|---|---|---|---|---|
| *lmo0136* | | 2.18 | **VirRS** [8] ↑ | Oligopeptide transport system permease protein | Amino acid transport and metabolism; Inorganic ion transport and metabolism |
| *lmo2433* | *estA* | 2.16 | **VirRS** [8] ↑ | Putative esterase | General function prediction only |
| *lmo2314* | | 2.09 | | Hypothetical protein | Not in COGs |
| *lmo2568* | | 2.06 | **CodY** [7] ↑ | Hypothetical protein | Not in COGs |
| *lmo2210* | | 2.04 | **LisR** [5] ↑ | Hypothetical protein | Not in COGs |
| *lmo1634* | *lap* | -7.69 | | Bifunctional acetaldehyde-CoA/alcohol dehydrogenase | Energy production and conversion |
| *lmo1591* | *argC* | -4.96 | **CodY** [7] ↓ | N-acetyl-gamma-glutamyl-phosphate reductase | Amino acid transport and metabolism |
| *lmo2090* | *argG* | -4.93 | **CodY** [7] ↓ | Argininosuccinate synthase | Amino acid transport and metabolism |
| *lmo2172* | | -4.28 | | Acetyl-CoA:acetoacetyl-CoA transferase, alpha subunit | Lipid transport and metabolism |
| *lmo1590* | *argJ* | -4.13 | **CodY** [7] ↓ | bifunctional ornithine acetyltransferase/N-acetylglutamate synthase protein | Amino acid transport and metabolism |
| *lmo1257* | | -4.04 | | Hypothetical protein | Not in COGs |
| *lmo2235* | | -3.91 | | Similar to NADH oxidase | Energy production and conversion; General function prediction only |
| *lmo2158* | | -3.59 | **SigB** [1] ↑ **CodY** [7] ↑ | hypothetical protein | Function unknown |
| *lmo2104* | *feoA* | -3.55 | **Fur** [4] ↓ | Hypothetical protein | Inorganic ion transport and metabolism |
| *lmo2091* | *argH* | -3.53 | **CodY** [7] ↓ | argininosuccinate lyase | Amino acid transport and metabolism |
| *lmo2234* | | -3.52 | **CodY** [7] ↓ | Inosose isomerase | Carbohydrate transport and metabolism |
| *lmo2250* | *arpJ* | -3.46 | **VirRS** [8] ↑ **CodY** [7] ↓ | Amino acid ABC transporter, amino acid-binding/permease protein | Amino acid transport and metabolism; Signal transduction mechanisms |
| *lmo2410* | | -3.45 | | Hypothetical protein | Not in COGs |
| *lmo2105* | *feoB* | -3.10 | **Fur** [4] ↓ | Ferrous iron transport protein B | Inorganic ion transport and metabolism |
| *rli119* | | -2.79 | | | |
| *lmo0412* | | -2.78 | | Hypothetical protein | Not in COGs |
| *lmo0654* | | -2.68 | **PrfA** [2] ↑; **CodY** [7] ↑ | Hypothetical protein | Not in COGs |
| *lmo2236* | | -2.65 | | Shikimate 5-dehydrogenase | Amino acid transport and metabolism |
| *lmo2173* | | -2.61 | **CodY** [7] ↑ | Sigma-54 dependent transcriptional regulator | Transcription; Signal transduction mechanisms |
| *lmo1406* | *pflB* | -2.60 | **CodY** [7] ↑ | pyruvate formate-lyase | Energy production and conversion |
| *lmo2251* | | -2.59 | **VirRS** [8] ↑ | Amino acid ABC transporter, ATP-binding protein | Amino acid transport and metabolism |
| *lmo2686* | *zea* | -2.55 | | Hypothetical protein | Not in COGs |

| | | | | | |
|---|---|---|---|---|---|
| *lmo1917* | *pflA* | -2.55 | | Pyruvate formate-lyase | Energy production and conversion |
| *lmo2238* | | -2.55 | | Major facilitator family transporter | Carbohydrate transport and metabolism; Amino acid transport and metabolism; Inorganic ion transport and metabolism; General function prediction only |
| *lmo2637* | *pplA* | -2.50 | | Putative pheromone precursor lipoprotein, related to Cad | Function unknown |
| *lmo0321* | | -2.48 | **CodY** [7] ↑ | membrane protein | Not in COGs |
| *lmo0912* | | -2.46 | **CodY** [7] ↓ | Formate efflux transporter | Inorganic ion transport and metabolism |
| *lmo0134* | | -2.40 | **SigB** [1] ↑; **PrfA** [2] ↑ | acetyltransferase, GNAT family | General function prediction only |
| *lmo0628* | | -2.40 | **CodY** [7] ↑ | Hypothetical protein | Not in COGs |
| *lmo0937* | | -2.37 | **PrfA** [2] ↑ | Hypothetical protein | Not in COGs |
| *lmo0515* | | -2.37 | **SigB** [1] ↑ | hypothetical protein | Signal transduction mechanisms |
| *lmo0133* | | -2.35 | **PrfA** [2] ↑ | hypothetical protein | Function unknown |
| *lmo2171* | | -2.30 | | Oxalate/formate antiporter | Carbohydrate transport and metabolism; Amino acid transport and metabolism; Inorganic ion transport and metabolism; General function prediction only; |
| *lmo0788* | | -2.28 | **PrfA** [2] ↑ | Activator of (R)-2-hydroxyglutaryl-CoA dehydratase | Lipid transport and metabolism; Function unknown |
| *lmo2067* | *bsh* | -2.23 | **SigB** [1] ↑; **PrfA** [2] ↑ | Choloylglycine hydrolase | Defense/virulence mechanisms |
| *lmo1131* | *cydC* | -2.22 | **Fur** [4] ↓ | Transport ATP-binding protein CydC | Energy production and conversion; Posttranslational modification, protein turnover, chaperones |
| *lmo2365* | | -2.22 | | Listeria RofA-like transcriptional regulator | Transcription; Carbohydrate transport and metabolism |
| *lmo1407* | *pflC* | -2.15 | | pyruvate-formate lyase activating enzyme | Posttranslational modification, protein turnover, chaperones |
| *lmo0302* | | -2.13 | | hypothetical protein | Not in COGs |
| *lmo2434* | *gadD* | -2.09 | **CodY** [7] ↑ | Glutamate decarboxylase | Amino acid transport and metabolism |
| *lmo2447* | | -2.08 | | transcriptional activator | Transcription |
| *rli105* | | -2.08 | | | |
| *lmo0953* | | -2.07 | **SigB** [1] ↑; **PrfA** [2] ↑; **CodY** [7] ↑ | hypothetical protein | Not in COGs |
| *lmo0309* | | -2.04 | | conserved hypothetical protein | Function unknown |
| *lmo2669* | | -2.02 | | membrane protein | Function unknown |

[a] Positive regulation (↑) represents higher transcript levels in the parent strain compared to the mutant strain in regulatory gene, whereas negative regulation (↓) represent higher transcript levels in the mutant strain in regulatory gene compared to the parent strain. The data are according to:
[1] Hain et al., 2008
[2] Milohanic et al., 2003

[3] van der Veen et al., 2010
[4] McLaughlin et al., 2012
[5] Nielsen et al., 2012
[6] Hu et al., 2007
[7] Bennett et al., 2007
[8] Mandin et al., 2005
[9] Rea et al., 2005
[b] Information from Listeriomics website (listeriomics.pasteur.fr)
