## Supplementary Table S5 for "Inactivation of *lmo0946* (*sif*) induces the SOS response and MGEs mobilization and silences the general stress response and virulence program in *Listeria monocytogenes*"

Table S5. Expression of MGE genes in *L. monocytogenes lmo0946**

| Gene name | Gene symbol | MGE name [1] | $log_2$ Fold Change | Padj[2] | Product [3] |
|---|---|---|---|---|---|
| ***lmo0113*** | | **monocin (lma operon): *lmo0113 - lmo0129*** | **0.98** | **8.38E-13** | similar to protein gp35 from Bacteriophage A118 |
| ***lmo0114*** | | | **0.92** | **1.44E-11** | similar to putative repressor C1 from lactococcal bacteriophage Tuc2009 |
| ***lmo0115*** | *lmaD* | | **3.74** | **1.63E-57** | Listeria protein LmaD. associated with virulence |
| ***lmo0116*** | *lmaC* | | **3.66** | **6.64E-60** | LmaC. associated with virulence in Listeria |
| ***lmo0117*** | *lmaB* | | **4.61** | **5.6E-134** | Antigen B |
| ***lmo0118*** | *lmaA* | | **4.52** | **4.3E-130** | Antigen A |
| ***lmo0119*** | | | **4.61** | **1.03E-65** | Hypothetical protein |
| ***lmo0120*** | | | **4.50** | **2.02E-84** | Hypothetical protein |
| ***lmo0121*** | | | **4.28** | **2.3E-109** | Phage tail length tape-measure protein |
| ***lmo0122*** | | | **4.23** | **1.6E-128** | Phage tail fiber |
| ***lmo0123*** | | | **4.15** | **1.3E-135** | Putative tail or base plate protein gp18 [Bacteriophage A118] |
| ***lmo0124*** | | | **3.96** | **9.22E-82** | Hypothetical protein |
| ***lmo0125*** | | | **4.26** | **5.7E-108** | Hypothetical protein |
| ***lmo0126*** | | | **4.32** | **4.82E-81** | Hypothetical protein |
| ***lmo0127*** | | | **4.24** | **8.1E-83** | Hypothetical protein |
| ***lmo0128*** | | | **3.99** | **5.77E-90** | Similar to phage-related protein |
| ***lmo0129*** | | | **4.12** | **2E-120** | N-acetylmuramoyl-L-alanine amidase |
| ***lmo1097*** | | **ICELm1 (Tn916): *lmo1097 - lmo1115*** | **13.71** | **4.63E-24** | Integrase. superantigen-encoding pathogenicity islands SaPI |
| *lmo1098* | | | 0.04 | NA | |
| *lmo1099* | | | 0.04 | NA | |
| ***lmo1100*** | *cadA* | | **12.27** | **3.03E-19** | Cadmium resistance protein |
| ***lmo1101*** | *lspB* | | **9.12** | **1.27E-10** | Hypothetical protein |
| ***lmo1102*** | *cadC* | | **7.85** | **7.51E-08** | Cadmium efflux system accessory protein |
| *lmo1103* | | | 0.15 | NA | |
| *lmo1104* | | | 0.22 | 0.249 | |
| ***lmo1105*** | | | **6.36** | **1.24E-04** | Membrane protein. putative |
| ***lmo1106*** | | | **5.18** | **0.002** | Hypothetical protein |
| *lmo1107* | | | 0.19 | NA | |
| *lmo1108* | | | 0.04 | NA | |
| *lmo1109* | | | 0.05 | NA | |
| *lmo1110* | | | 0.00 | NA | |
| *lmo1111* | | | 0.32 | 0.138 | |
| ***lmo1112*** | | | **5.29** | **0.001** | Hypothetical protein |
| *lmo1113* | | | 4.35 | 0.010 | |
| *lmo1114* | | | 0.46 | 0.064 | |
| ***lmo1115*** | | | **4.96** | **0.002** | Similar to fibrinogen-binding protein |
| ***lmo2271*** | | **A118: *lmo2271 - lmo2332*** | **3.51** | **1.82E-42** | Hypothetical protein |
| *lmo2272* | | | 0.20 | 0.657 | |
| *lmo2273* | | | 0.13 | 0.565 | |
| *lmo2274* | | | 0.62 | 0.012 | |
| ***lmo2275*** | | | **0.73** | **2.97E-05** | Portein gp28 [Bacteriophage A118] |
| *lmo2276* | | | 0.04 | 0.855 | |
| *lmo2277* | | | 0.50 | 0.013 | |
| ***lmo2278*** | *lysA* | | **3.09** | **4.3E-20** | L-alanoyl-D-glutamate peptidase |
| ***lmo2279*** | | | **2.78** | **4E-08** | Holin |
| ***lmo2280*** | | | **2.45** | **1.50E-04** | Protein gp23 |
| *lmo2281* | | | 0.04 | NA | |
| ***lmo2282*** | | | **3.59** | **6.92E-16** | protein gp21 |
| ***lmo2283*** | | | **2.84** | **3.26E-21** | Protein gp20 |
| ***lmo2284*** | | | **3.13** | **1.65E-39** | Protein gp19 |

| | | | | |
|---|---|---|---|---|
| ***lmo2285*** | | **3.51** | **3.8E-47** | Protein gp18 |
| ***lmo2286*** | | **2.75** | **1.46E-22** | Protein gp17 |
| ***lmo2287*** | | **3.20** | **3.91E-70** | Putative tape-measure |
| ***lmo2288*** | | **3.74** | **6.22E-28** | Protein gp15 |
| ***lmo2289*** | | **4.35** | **4.71E-26** | Protein gp14 |
| ***lmo2290*** | | **3.24** | **1E-07** | Protein gp13 |
| ***lmo2291*** | | **3.68** | **2.36E-45** | Major tail shaft protein |
| ***lmo2292*** | | **4.19** | **1.47E-40** | Protein gp11 |
| ***lmo2293*** | | **3.39** | **6.08E-45** | Protein gp10 |
| ***lmo2294*** | | **4.53** | **9.8E-35** | Protein gp9 |
| ***lmo2295*** | | **3.30** | **2.07E-42** | Protein gp8 |
| ***lmo2296*** | | **3.62** | **1.66E-94** | Phage capsid protein |
| ***lmo2297*** | | **3.29** | **2.51E-35** | Putative scaffolding protein |
| ***lmo2298*** | | **3.81** | **1.68E-91** | Protein gp4 |
| ***lmo2299*** | | **3.46** | **2.4E-106** | Putative portal protein |
| ***lmo2300*** | | **3.33** | **9.31E-43** | Putative terminase large subunit from bacteriophage A118 |
| ***lmo2301*** | | **3.14** | **3.03E-21** | Terminase small subunit [Bacteriophage A118] |
| ***lmo2302*** | | **0.81** | **3.24E-04** | Hypothetical protein |
| ***lmo2303*** | | **3.06** | **4.53E-29** | Protein gp66 |
| *lmo2304* | | 0.22 | 0.273 | |
| ***lmo2305*** | | **2.80** | **2.02E-23** | Hypothetical protein. Lmo2305 homolog [Bacteriophage A118] |
| ***lmo2306*** | | **2.34** | **6.51E-10** | Hypothetical protein |
| *lmo2307* | | 0.55 | 0.241 | |
| ***lmo2308*** | | **2.34** | **7.28E-16** | Single-stranded DNA-binding protein (prophage associated) |
| *lmo2309* | | 0.89 | 0.074 | |
| *lmo2310* | | 0.39 | 0.189 | |
| ***lmo2311*** | | **2.46** | **3.17E-06** | Hypothetical protein |
| ***lmo2312*** | | **2.79** | **1.04E-06** | Conserved hypothetical protein |
| ***lmo2313*** | | **3.75** | **6.7E-29** | Hypothetical protein. Lmo2313 homolog [Bacteriophage A118] |
| ***lmo2314*** | | **3.88** | **0.003** | Hypothetical protein |
| ***lmo2315*** | | **3.90** | **3.23E-41** | Protein gp51 [Bacteriophage A118] |
| ***lmo2316*** | | **3.88** | **2.46E-50** | Methyltransferase |
| ***lmo2317*** | | **3.35** | **4.99E-35** | Protein gp49. replication initiation [Bacteriophage A118] |
| ***lmo2318*** | | **3.52** | **1.06E-23** | Putative recombination protein / Single-stranded DNA-binding protein |
| ***lmo2319*** | | **3.50** | **1.29E-24** | Hypothetical protein |
| ***lmo2320*** | | **2.88** | **2.24E-04** | Hypothetical protein |
| ***rli140*** | | **3.14** | **2.66E-04** | |
| ***lmo2321*** | | **4.35** | **2.42E-12** | Protein gp45 [Bacteriophage A118] |
| ***lmo2322*** | | **3.64** | **1.26E-15** | gp44 |
| ***lmo2323*** | | **3.58** | **3.78E-38** | Protein gp43 [Bacteriophage A118] |
| ***lmo2324*** | | **4.12** | **2.81E-59** | Phage antirepressor protein / Antirepressor [Bacteriophage A118] |
| ***lmo2325*** | | **4.81** | **2.65E-21** | Hypothetical protein |
| ***lmo2326*** | | **3.82** | **1.89E-11** | Protein gp41 [Bacteriophage A118] |
| ***lmo2327*** | | **3.59** | **3.08E-22** | Hypothetical protein |
| ***lmo2328*** | | **1.84** | **0.009554** | Similar to transcription regulator |
| *lmo2329* | | 0.03 | 0.887 | |
| ***lmo2330*** | | **0.49** | **0.004** | Similar to protein gp33 [Bacteriophage A118] |
| *lmo2331* | | 0.08 | 0.781 | |
| ***int*** | | **1.25** | **6.03E-16** | Integrase [Bacteriophage A118] |
| *lmo0459* | **IS3-I: *lmo0459 - lmo463*** | 0.40 | 0.034 | |
| *lmo0460* | | 0.18 | 0.612 | |
| *lmo0461* | | -0.14 | 0.707 | |

| | | |
|---|---|---|
| *lmo0462* | -0.14 | 0.756 |
| *lmo0463* | -0.02 | 0.962 |

[1] Information on MGE according to Kuenne et al.. 2013 (Kuenne C. Billion A. Mraheil MA. Strittmatter A. Daniel R. Goesmann A. Barbuddhe S. Hain T. Chakraborty T. Reassessment of the *Listeria monocytogenes* pan-genome reveals dynamic integration hotspots and mobile genetic elements as major components of the accessory genome. BMC Genomics. 2013. 22;14:47. doi: 10.1186/1471-2164-14-47)

[2] $Log_2$ expression levels with an adjusted p value of MGE genes in mutant *lmo0946** vs *Listeria monocytogenes EGD-e* from exponential phase of growth in BHI in 37 °C; In bold genes with upregulated expression (Padj < 0.01)

[3] Information on upregulated MGE genes from Listeriomics website (listeriomics.pasteur.fr)
