## Supplementary Table S6 for "Inactivation of *lmo0946* (*sif*) induces the SOS response and MGEs mobilization and silences the general stress response and virulence program in *Listeria monocytogenes*"

**Table S6. Comparison of expression of SOS response genes in *L. monocytogenes lmo0946** to the expression observed in wild-type *L. monocytogenes* EGD-e strain after MMC treatment.**

| Gene name | Gene symbol | Description of product[1] | MMC treatment wt[2] | *lmo0946**[3] | Padj[3] |
|---|---|---|---|---|---|
| ***lmo0157*** | | Predicted ATP-dependent helicase | 1.61 | **1.06** | **1.97E-21** |
| *lmo0158* | | Predicted hydrolase | 1.18 | 0.13 | 0.53 |
| ***lmo1302*** | *lexA* | Transcription repressor of SOS response | 1.74 | **0.77** | **9.71E-09** |
| ***lmo1303*** | *yneA* | Similar to B. subtilis YneA protein | 3.95 | **1.91** | **0.001** |
| ***lmo1398*** | *recA* | Transcription activator of SOS response | 2.58 | **1.23** | **1.33E-38** |
| *lmo1421* | *bilEA* | Osmoprotectant transport system ATP-binding protein, bile resistance | 1.13 | -0.07 | 0.81 |
| *lmo1422* | *bilEB* | Osmoprotectant transport system permease protein, bile resistance | 1.21 | 0.09 | 0.74 |
| ***lmo1574*** | *dnaE* | DNA polymerase III alpha subunit | 1.37 | **0.61** | **1.52E-10** |
| ***lmo1640*** | | Hypothetical protein | 1.90 | **1.61** | **1.2E-12** |
| ***lmo1639*** | | DNA-3-methyladenine glycosidase, base excision repair | 2.24 | **1.89** | **1.8E-40** |
| ***lmo1638*** | | redicted peptidase | 2.22 | **1.62** | **1.08E-25** |
| ***lmo1759*** | *pcrA* | ATP-dependent DNA helicase | 0.93 | **0.50** | **1.49E-08** |
| ***lmo1758*** | *ligA* | NAD-dependent DNA ligase | 0.65 | **0.42** | **3.36E-05** |
| ***lmo1975*** | *dinB* | DNA polymerase IV | 3.08 | **2.29** | **6.85E-96** |
| ***lmo2222*** | | Predicted DNA repair exonuclease | 1.95 | **1.54** | **1.8E-38** |
| ***lmo2221*** | | Hypothetical protein | 2.38 | **1.53** | **2.93E-46** |
| ***lmo2220*** | | Predicted exonuclease | 1.52 | **0.74** | **6.64E-18** |
| ***lmo2268*** | *addB* | Predicted ATP-dependent helicase | 1.59 | **1.05** | **4.96E-31** |
| ***lmo2267*** | | Predicted ATP-dependent helicase | 1.74 | **0.87** | **2.69E-18** |
| *lmo2266* | | Predicted hydrolase | 1.56 | 0.44 | 0.12 |
| ***lmo2265*** | | Hypothetical protein | 1.61 | **1.32** | **5.05E-08** |
| ***lmo2264*** | | Hypothetical protein | 1.28 | **0.81** | **2.98E-12** |
| ***lmo2271*** | | Bacteriophage A118 protein | 3.4 | **3.51** | **1.82E-42** |
| ***lmo2332*** | *int* | Site-specific DNA recombinase, integrase (Bacteriophage A118) | 1.58 | **1.25** | **6.03E-16** |
| ***lmo2489*** | *uvrB* | Excinuclease ABC (subunit B) | 2.76 | **1.82** | **1.8E-34** |
| ***lmo2488*** | *uvrA* | Excinuclease ABC (subunit A) | 2.63 | **1.61** | **1.75E-58** |
| ***lmo2675*** | *umuD* | DNA polymerase V | 2.93 | **3.70** | **3E-128** |
| ***lmo2676*** | *umuC* | DNA polymerase V | 2.14 | **3.70** | **1.8E-172** |
| ***lmo2828*** | | Predicted equivalent to the UmuD subunit of polymerase V from Gram-negative bacteria | 4.56 | **2.58** | **8.74E-17** |

[1] Information from van der Veen et al., 2010

[2] $Log_2$ expression levels of genes from SOS regulon in the wild-type (Wt) strain after MMC treatment vs the wild-type untreated according to van der Veen et al., 2010 (van der Veen S., van Schalkwijk S., Molenaar D., de Vos W. M., Abee T., Wells-Bennik M. H. J. (2010). The SOS response of *Listeria monocytogenes* is involved in stress resistance and mutagenesis. Microbiology 156, 374-384. DOI 10.1099/mic.0.035196-0.)

[3] $Log_2$ expression levels of genes from SOS regulon in mutant *lmo0946** vs *Listeria monocytogenes EGD-e* from exponential phase of growth without stress factors; in bold genes with essentially changed expression (Padj<0.01)
