## Supplementary Table S7 for "Inactivation of *lmo0946* (*sif*) induces the SOS response and MGEs mobilization and silences the general stress response and virulence program in *Listeria monocytogenes*"

Table S7. Expression of PrfA regulon in *L. monocytogenes lmo0946**

| Gene Group[1] | Gene name[1] | Regulation[1] | *lmo0946**[2] | Padj[2] | Description of product[1] |
|---|---|---|---|---|---|
| **Group I** (positively regulated and preceeded by PrfA box; downregulated in presence of celobiose and upregulated in presence of charcoal) | ***hly*** | PrfA box | **-0.49** | **0.001** | listeriolysin O precursor |
| | ***mpl*** | PrfA box | **-0.66** | **0.019** | Zinc metalloproteinase precursor |
| | ***actA*** | PrfA box | **-0.52** | **0.014** | actin-assembly inducing protein precursor |
| | *plcB* | PrfA box | -0.09 | 0.756 | |
| | *plcA* | PrfA box | -0.46 | 0.126 | |
| | ***prfA*** | PrfA box; SigB | **1.36** | **7.79E-10** | Listeriolysin O positive regulatory protein |
| | ***inlA*** | PrfA box | **-1.30** | **3.02E-05** | Internalin A |
| | ***inlB*** | PrfA box | **-1.09** | **4.93E-04** | Internalin B |
| | *inlC* | PrfA box | 0.40 | 0.167 | |
| | *uhpT* | PrfA box | -0.45 | 0.262 | |
| | *lmo2219* | PrfA box | -0.18 | 0.36 | |
| | ***lmo0788*** | PrfA box | **-2.28** | **1.30E-38** | Unknown |
| **Group II** (negatively regulated by PrfA) | *lmo0178* | | -0.42 | 0.149 | |
| | *lmo0179* | | -0.05 | 0.860 | |
| | *lmo0180* | | -0.16 | 0.527 | |
| | *lmo0181* | | 0.19 | 0.431 | |
| | *lmo0182* | | 0.23 | 0.304 | |
| | *lmo0183* | | 0.31 | 0.058 | |
| | *lmo0184* | | -0.02 | 0.931 | |
| | ***lmo0278*** | PrfA box | **-0.83** | **1.49E-05** | Unknown, similar to sugar ABC transporter, ATP-binding protein |
| **Group III** (positively regulated by PrfA; upregulated in presence of celobiose and downregulated in presence of charcoal) | ***lmo0596*** | PrfA box; SigB | **-1.09** | **0.0367** | Unknown, similar to unknown proteins |
| | ***lmo2067*** | PrfA box; SigB | **-2.23** | **2.74E-08** | Unknown, similar to conjugated bile acid hydrolase |
| | *opuCA* | SigB | -0.51 | 0.268 | |
| | *opuCB* | | -0.49 | 0.285 | |
| | *opuCC* | | -0.35 | 0.430 | |
| | *opuCD* | | -0.52 | 0.258 | |
| | ***lmo1602*** | SigB | **-0.71** | **1.12E-08** | similar to general stress protein |
| | ***lmo1601*** | | **-0.66** | **2.29E-08** | similar to general stress protein |
| | ***lmo2748*** | SigB | **-1.25** | **8.12E-04** | Unknown, similar to *B. subtilis* stress protein YdaG |
| | *lmo2230* | SigB | -0.53 | 0.237 | |
| | *lmo2231* | | -0.39 | 0.364 | |
| | *lmo0913* | SigB | -0.89 | 0.094 | |
| | *lmo0669* | SigB | -0.52 | 0.153 | |
| | ***lmo0670*** | | **-0.90** | **0.023** | Unknown |
| | ***lmo2573*** | | **-1.34** | **0.013** | Unknown, similar to zinc-binding dehydrogenase |
| | ***lmo2572*** | | **-1.13** | **0.004** | Unknown, similar to Chain A, Dihydrofolate Reductase |
| | ***lmo2571*** | | **-1.38** | **6.82E-05** | Unknown, similar to nicotinamidase |
| | *lmo2570* | | -0.32 | 0.471 | |
| | *lmo2695* | SigB | -0.82 | 0.095 | |
| | *lmo2696* | | -0.47 | 0.297 | |
| | *lmo2697* | | -0.39 | 0.391 | |
| | *lmo1694* | SigB | -0.86 | 0.101 | |
| | *lmo0539* | SigB | -0.66 | 0.160 | |
| | ***lmo0784*** | SigB | **-1.79** | **1.07E-11** | Unknown, similar to mannose-specific phosphotransferase system (PTS) component IIA |
| | ***lmo0783*** | | **-1.43** | **2.39E-05** | Unknown, similar to mannose-specific phosphotransferase system (PTS) component IIB |
| | ***lmo0782*** | | **-1.00** | **0.005** | Unknown, similar to mannose-specific phosphotransferase system (PTS) component IIC |
| | *lmo0781* | | -0.49 | 0.278 | |

| | | | | |
|---|---|---|---|---|
| ***lmo0602*** | SigB | **-1.56** | **4.05E-06** | Unknown, weakly similar to transcription regulator |
| ***lmo2391*** | SigB | **-1.39** | **4.31E-05** | Unknown, conserved hypothetical protein similar to *B. subtilis* YhfK protein |
| ***lmo0043*** | | **-1.84** | **4.79E-07** | Unknown, similar to arginine deiminase |
| ***lmo0133*** | | **-2.35** | **4.34E-16** | Unknown, similar to *E. coli* YjdI protein |
| ***lmo0134*** | | **-2.40** | **8.36E-16** | Unknown, similar to *E. coli* YjdJ protein |
| ***lmo0937*** | SigB | **-2.37** | **3.70E-15** | Unknown |
| *lmo0994* | SigB | -0.65 | 0.174 | |
| ***lmo0794*** | SigB | **-1.05** | **0.004** | Unknown, similar to *B. subtilis* YwnB protein |
| ***lmo0654*** | | **-2.68** | **7.63E-33** | Unknown |
| ***lmo2213*** | SigB | **-1.47** | **0.025** | Unknown, similar to unknown protein |
| ***lmo2673*** | | **-1.43** | **0.016** | Unknown, conserved hypothetical protein |
| ***sepA*** | SigB | **-1.11** | **0.047** | Unknown |
| ***lmo0796*** | | **-1.02** | **1.29E-05** | Unknown, conserved hypothetical protein |
| ***lmo1261*** | SigB | **-0.91** | **0.010** | Unknown |
| ***lmo0439*** | SigB | **-1.35** | **0.018** | Unknown, weakly similar to a module of peptide synthetase |
| ***lmo0953*** | | **-2.07** | **6.20E-14** | Unknown |
| ***lmo0019*** | | **-1.09** | **0.017** | Unknown |
| *lmo0555* | | -0.54 | 0.058 | |
| *lmo0676* | | 0.45 | 0.215 | |
| *inlH* | | -0.87 | 0.102 | |
| ***lmo0169*** | | **-1.41** | **1.59E-04** | Unknown, similar to a glucose uptake protein |
| ***lmo0170*** | | **-0.71** | **0.020** | Unknown |
| *lmo0242* | | -0.26 | 0.167 | |
| ***rsbV*** | SigB | **-1.05** | **1.47E-06** | anti-anti-sigma factor (antagonist of RsbW) |
| ***lmo0641*** | SigB | **1.14** | **2.59E-07** | Unknown, similar to heavy metal-transporting ATPase |
| ***galE*** | | **-0.34** | **0.006** | UDP-glucose 4-epimerase |

[1] Information from Milohanic et al., 2003 (Milohanic, E., Glaser, P., Coppee, J.Y., Frangeul, L., Vega, Y., Vazquez-Boland, J.A., et al. (2003). Transcriptome analysis of *Listeria monocytogenes* identifies three groups of genes differently regulated by PrfA. Mol Microbiol 47(6), 1613-1625.); genes in operons are boxed

[2] $Log_2$ expression levels with an adjusted p value of genes from PrfA regulon in mutant *lmo0946** vs *Listeria monocytogenes EGD-e* from exponential phase of growth without stress factors;

In red genes with upregulated expression (Padj<0.01); in dark blue genes with downregulated expression (Padj<0.01); in light blue genes with downregulated expression (Padj<0.05);
