## Supplementary Table S8 for "Inactivation of *lmo0946* (*sif*) induces the SOS response and MGEs mobilization and silences the general stress response and virulence program in *Listeria monocytogenes*"

Table S8. Expression levels of genes from *sigB* operon in mutant *lmo0946** vs *Listeria monocytogenes EGD-e* from exponential phase of growth in 37 °C.

| Gene name | Gene symbol | $log_2$ Fold Change | Padj | Product [1] |
|---|---|---|---|---|
| *lmo0889* | *RsbR* | 0.039 | 0.758 | RsbR, positive regulator of sigma-B |
| *lmo0890* | *rsbS* | 0.058 | 0.728 | RsbS, negative regulator of sigma-B |
| *lmo0891* | *rsbT* | 0.228 | 0.103 | Anti-sigma B factor RsbT |
| *lmo0892* | *rsbU* | -0.061 | 0.72 | Serine phosphatase RsbU, regulator of sigma subunit |
| *lmo0893* | *rsbV* | -1.048 | 1.47E-06 | anti-anti-sigma factor (antagonist of RsbW) |
| *lmo0894* | *rsbW* | -0.478 | 0.002 | serine-protein kinase RsbW |
| *lmo0895* | *sigB* | -0.355 | 0.138 | RNA polymerase sigma factor SigB |
| *lmo0896* | *rsbX* | -0.281 | 0.193 | indirect negative regulation of sigma B dependant gene expression (serine phosphatase) |

[1] Information from Listeriomics website (listeriomics.pasteur.fr)
Genes precceded by SigB promoter located within the *sigB* operon are boxed.
