## Supplementary Table S9 for "Inactivation of *lmo0946* (*sif*) induces the SOS response and MGEs mobilization and silences the general stress response and virulence program in *Listeria monocytogenes*"

**Supplementary Table S9:** Primers used in this study

| Name | Sequence (5’→ 3’) | Further information |
|---|---|---|
| | **Mutants construction** | |
| **DL0944FA** | ATGGATCCTGCGAGCGTAGAAGAAGCG | Forward primer for upstream flanking region of *lmo0944*. EcoRI restriction enzyme site (underlined). |
| **DL0944RB** | TCATGCTTGGTCCGTTACCTC | Reverse primer for upstream flanking region of *lmo0944*. |
| **DL0944FC** | **GTAACGGACCAAGCATGA**AGCAGCTGTAGCTGAGAAG | Forward primer for downstream flanking region of *lmo0944*. Anneals with DL0944RB (bold). |
| **DL0944RD** | TGTCCATGGTCTGCGTGCGGGTGTG | Reverse primer for downstream flanking region of *lmo0944*. NcoI restriction enzyme site (underlined). |
| **DL0945FA** | GTGGATCCTGTGCGGGATTAATCGTCTC | Forward primer for upstream flanking region of *lmo0945*. EcoRI restriction enzyme site (underlined). |
| **DL0945RB** | CTATGTCATCTTTCTCCTCCTGC | Reverse primer for upstream flanking region of *lmo0945*. |
| **DL0945FC** | **AGGAGAAAGATGACATAG**ACGCGGCGAAAAAGCGA | Forward primer for downstream flanking region of *lmo0945*. Anneals with DL0945RB (bold). |
| **DL0945RD** | GTGCCATGGTCATCTATGGACTGATATGGT | Reverse primer for downstream flanking region of *lmo0945*. NcoI restriction enzyme site (underlined). |
| **Fri Fpm** | GTGGATCCGCTTGGGCCTCCTAGTTG | Forward primer for region of *fri*. EcoRI restriction enzyme site (underlined). |
| **Fri Rpm** | GTTCCATGGCTTGGTCCGTTACCTCG | Reverse primer for region of *fri*. NcoI restriction enzyme site (underlined). |
| **Fri F sub** | CAAGTAGCGAATT**AG**AACGTATTCACA | Forward primer for site-directed mutagenesis of *fri*. Substituted nucleotides (bold). |
| **Fri R sub** | TGTGAATACGTT**CT**AATTCGCTACTTG | Reverse primer for site-directed mutagenesis of *fri*. Substituted nucleotides (bold). |
| **Fri F subsc** | GTTATCGCCGCATTTGTTCT**AG** | Forward primer for verification of chromosomal substitution in *fri*, used in pair with Fri Rpm. Substituted nucleotides (bold). |
| **Lmo0946 Fpm** | CAGGATCCGAAGAATTACCAGAAAGC | Forward primer for region of *lmo0946* and *lhrC5*. EcoRI restriction enzyme site (underlined). |
| **LhrC5 Rpm** | CTTCCATGGCACAATCTATCGGTTACT | Reverse primer for region of *lmo0946* and *lhrC5*. NcoI restriction enzyme site (underlined). |

| | | |
|---|---|---|
| **0946 F sub** | CAGACCAAACA**T**A**G**TGGCAAATTGAC | Forward primer for site-directed mutagenesis of *lmo0946*. Substituted nucleotides (bold). |
| **0946 R sub** | GTCAATTTGCCA**C**T**A**TGTTTGGTCTG | Reverse primer for site-directed mutagenesis of *lmo0946*. Substituted nucleotides (bold). |
| **0946 F subsc** | CTATTAGAACCAGACCAAACA**T**A**G** | Forward primer for verification of chromosomal substitution in *lmo0946*, used in pair with LhrC5 Rpm. Substituted nucleotides (bold). |
| **LhrC5 F sub** | AGAAAAGT**C**A**G**T**GC**ATGTATAAGCTAACAACA | Forward primer for site-directed mutagenesis of *lhrC5*. Substituted nucleotides (bold). |
| **LhrC5 R sub** | TGTTGTTAGCTTATACAT**GC**A**C**T**G**ACTTTTCT | Reverse primer for site-directed mutagenesis of *lhrC5*. Substituted nucleotides (bold). |
| **LhrC5 R subsc** | GTTGTTAGCTTATACAT**GC**A**C**T**G** | Reverse primer for verification of chromosomal substitution in *lhrC5*, used in pair with Lmo0946 Fpm. Substituted nucleotides (bold). |
| **LhrC1-4FA** | CGAGTCGACACCGGGACCAGATGGCATG | Forward primer for upstream flanking region of *lhrC1*. SalI restriction enzyme site (underlined). |
| **LhrC1-4RB** | CCAAGAAAAGAAAGTTTCTGCTGTGC | Reverse primer for upstream flanking region of *lhrC1*. |
| **LhrC1-4FC** | **CACAGCAGAAACTTTCTTTTCTTGG**GTGCGCTAAAATC ACTAAATTCTGC | Forward primer for downstream flanking region of *lhrC4*. Anneals with LhrC1-4RB (bold). |
| **LhrC1-4RD** | CGAAGATCTGCTCGTCTGGTACTGATGCTGC | Reverse primer for downstream flanking region of *lhrC4*. BglII restriction enzyme site (underlined). |
| **Rli22FA** | AGAGTCGACGAAGTTATTGGAGGAAACATTCG | Forward primer for upstream flanking region of *rli22* (*lhrC6*). SalI restriction enzyme site (underlined). |
| **Rli22RB** | AAGTTCTATATTAAAGGTTAATTATGAAC | Reverse primer for upstream flanking region of *rli22* (*lhrC6*). |
| **Rli22FC** | **GTTCATAATTAACCTTTAATATAGAACTT**GGACCGACT TGATTGTCGG | Forward primer for downstream flanking region of *rli22* (*lhrC6*). Anneals with Rli22RB (bold). |
| **Rli22RD** | AGAAGATCTGCAATTAGTCCGCCAACAG | Reverse primer for downstream flanking region of *rli22* (*lhrC6*). BglII restriction enzyme site (underlined). |
| **Rli33-1FA** | ACAGTCGACGGACGACAACTTAGAAAAAG | Forward primer for upstream flanking region of *rli33-1* (*lhrC7*). SalI restriction enzyme site (underlined). |
| **Rli33-1RB** | GTTATATAATACCCGTCTCAC | Reverse primer for upstream flanking region of *rli33-1* (*lhrC7*)). |
| **Rli33-1FC** | **GTGAGACGGGTATTATATAAC**GGAAAAAATGCTAAATA TAAGAAAACTCC | Forward primer for downstream flanking region of *rli33-1* (*lhrC7*). Anneals with Rli33-1RB (bold). |

| | | |
|---|---|---|
| **Rli33-1RD** | ACAAGATCTTGGTAATAACATACACTGCTAAAAC | Reverse primer for downstream flanking region of *rli33-1* (*lhrC7*). BglII restriction enzyme site (underlined). |
| **Lmo0946 Fcom** | CTTCTGCAGCATTGAAGATCATGATTATTGGTATTTC | Forward primer for complementation of *lmo0946**. PstI restriction enzyme site (underlined). |
| **Lmo0946 Rcom** | CTTGGTACCGGGAGTTTGGGATTTCCATTC | Reverse primer for complementation of *lmo0946**. KpnI restriction enzyme site (underlined). |
| **RT-PCR** | | |
| **RTLhrC5** | AGGGAGTAAACCGCACTAG | Primer used for reverse transcription in cotranscription analysis |
| **0943F** | CATTGGTATATGAGAGGCCAC | Forward primer used for PCR of *fri* in cotranscription analysis |
| **0943R** | CATTGTCGCCTTCTTTGTCAG | Reverse primer used for PCR of *fri* in cotranscription analysis |
| **0944F** | ATGGTTTCATGATGAGTTTGATGT | Forward primer used for PCR of *lmo0944* in cotranscription analysis |
| **0944R** | TCCGTTTTTGGTTCATAGTCG | Reverse primer used for PCR of *lmo0944* in cotranscription analysis |
| **lmaA F** | CAAGGTCTAACTGTAAACCGTTCT | Forward primer used for RT-qPCR of *lmaA* |
| **lmaA R** | CCAATTCTTTGTGAGACTCTGCATC | Reverse primer used for RT-qPCR of *lmaA* |
| **lmo2676 F** | GCCAAACTCGCGCTCGATAATG | Forward primer used for RT-qPCR of *lmo2676* |
| **lmo2676 R** | GAACAGCAGTCCGACGTCCAATT | Reverse primer used for RT-qPCR of *lmo2676* |
| **lmo2308 F** | CTGAGGTAGTTGCTGAATCAGTTCAATT | Forward primer used for RT-qPCR of *lmo2308* |
| **lmo2308 R** | ATCGCTCTTCTGACTCGTATCCGC | Reverse primer used for RT-qPCR of *lmo2308* |
| **sigB F** | ACTTCAAAGCTCGCCGCAAATTAG | Forward primer used for RT-qPCR of *sigB* |
| **sigB R** | ATCGTACTTCCATCCGAATCAGCTT | Reverse primer used for RT-qPCR of *sigB* |
| **rsbV F** | GACATATTTGTTGCTGGGGAGATCG | Forward primer used for RT-qPCR of *rsbV* |
| **rsbV R** | TACAAATACGCCTAATCCGGTGCTATC | Reverse primer used for RT-qPCR of *rsbV* |
| **lmo1634 F** | ATCAGTAGAGTGAATAACTGCGG | Forward primer used for RT-qPCR of *lmo1634* |
| **lmo1634 R** | GTGTTGGCGACAAATACCCA | Reverse primer used for RT-qPCR of *lmo1634* |
| **int F** | GCAATGGATAGAGCAATCGTTTTAGG | Forward primer used for RT-qPCR of *int* |
| **int R** | CTTCACCGATTCTCATGCCTGTC | Reverse primer used for RT-qPCR of *int* |
| **uvrA F** | TGGAAACACGCTTATTGTCGTTGAG | Forward primer used for RT-qPCR of *uvrA* |
| **uvrA R** | ACGTTTAGCAGGGACTGGAA | Reverse primer used for RT-qPCR of *uvrA* |
| **prfA F** | GCAGGCTACCGCATACGTTATC | Forward primer used for RT-qPCR of *prfA* |
| **prfA R** | TTCTTTACCATACACATAGGTCAGGA | Reverse primer used for RT-qPCR of *prfA* |
| **zea F** | GGGAATATAGCTCAAGAGAAAGAAATTG | Forward primer used for RT-qPCR of *zea* |
| **zea R** | ACCCTTACTAACATTAATCTTAGCATTAAACGA | Reverse primer used for RT-qPCR of *zea* |
| **hfq F** | GGTGGACAAGGGTTACAGGA | Forward primer used for RT-qPCR of *hfq* |

| | | |
|---|---|---|
| **hfq R** | ACAACGCGTCCTCTTAACTGA | Reverse primer used for RT-qPCR of *hfq* |
| **lmo2105 F** | GCTCGTCTTTGTTCCTCAAATCTTG | Forward primer used for RT-qPCR of *lmo2105* |
| **lmo2105 R** | CAATCGACCGAGCTGCCATAATTC | Reverse primer used for RT-qPCR of *lmo2105* |
| **argG F** | TCTCCTGTCCGTGATTGGAAATG | Forward primer used for RT-qPCR of *agrG* |
| **argG R** | GCACACCACATTCATTACTTCTAC | Reverse primer used for RT-qPCR of *agrG* |
| **lmo2279 F** | CCGCCTAAGTGGCTTCCGAC | Forward primer used for RT-qPCR of *lmo2279* |
| **lmo2279 R** | TTGCAAGCGATCCAGAGCCG | Reverse primer used for RT-qPCR of *lmo2279* |
| **recA F** | GCTCATGTTGGATTACAAGCACG | Forward primer used for RT-qPCR of *recA* |
| **recA R** | ACGTACAGTAGAATAGAATTTAAGCGCA | Reverse primer used for RT-qPCR of *recA* |
| **hly F** | CCTCCTGCATATATCTCAAGTGTG | Forward primer used for RT-qPCR of *hly* |
| **hly R** | GAACCTCCGTAAATTACGGCTTTGA | Reverse primer used for RT-qPCR of *hly* |
| **rpoB F** | CGTCGTCTTCGTTCTGTTGG | Forward primer used for RT-qPCR of *rpoB* |
| **rpoB R** | GTTCACGAACCACACGTTCC | Reverse primer used for RT-qPCR of *rpoB* |
| | **Northern blot analysis** | |
| **Fri NB F** | GGCGAACAAATGGATGAAGTA | Forward primer for double stranded probe for *fri* mRNA |
| **Fri NB R** | CAATACCTTGTTGATATTCGTC | Reverse primer for double stranded probe for *fri* mRNA |
| **lmo0944 NB F** | GGTAATGGAATTCGACTTTTTGC | Forward primer for double stranded probe for *lmo0944* mRNA |
| **lmo0944 NB R** | GGTTCATAGTCGATTTTCCAG | Reverse primer for double stranded probe for *lmo0944* mRNA |
| **lmo0945 NB F** | CCAAGACGCGGCGAAAAAGC | Forward primer for double stranded probe for *lmo0945* mRNA |
| **lmo0945 NB R** | CGCTTCTAAATTGGCACCGC | Reverse primer for double stranded probe for *lmo0945* mRNA |
| **lmo0946 NB F** | ATGAAAAAAGCAATTTTAGATCGG | Forward primer for double stranded probe for *sif* mRNA |
| **lmo0946 NB R** | GATGAATCGAGCTTTTCTTTG | Reverse primer for double stranded probe for *sif* mRNA |
| **lhrC5 NB F** | ATAAGCTAACAACAAGCAAAACATTTTCATTTCTTTCCC TTTTTA**GAATGGAAATCCCAAACTCCC** | Forward primer for double stranded probe for *lhrC5* mRNA. Anneals with lhrC5 NB R (bold). |
| **lhrC5 NB R** | AAAAAAACTAGTGCGGAAAAAGGGAGTAAACCGCACTA GCTAAAA**GGGAGTTTGGGATTTCCATTC** | Reverse primer for double stranded probe for *lhrC5* mRNA. Anneals with lhrC5 NB F (bold). |
| | **MGE analysis** | |
| **A118 R 1** | TGTATCACTTGAACGCTTTGAC | Reverse primer no 1 for A118 mobilization analysis |
| **A118 F 2** | TTAGCTGATTTAGCAACAGTTGAT | Forward primer no 2 for A118 mobilization analysis |
| **A118 F 3** | ATG AAA AAA GAA CAA ATC AGT ACT CAG | Forward primer no 3 for A118 mobilization analysis |
| **A118 R 4** | TTA TTG CTC GGG ATC TTG AGG AT | Reverse primer no 4 for A118 mobilization analysis |
| **ICELm1 R 1** | GAATTATTTCCTTTCAACCCGCTGAGC | Reverse primer no 1 for ICElm1 mobilization analysis |
| **ICELm1 F 2** | GCTGTAGGTTCATTCAACACAAGA | Forward primer no 2 for ICElm1 mobilization analysis |

| | | |
|---|---|---|
| **ICELm1 F 3** | GTTATGGGTGATGGTAGAACGTATGACC | Forward primer no 3 for ICElm1 mobilization analysis |
| **ICELm1 R 4** | AGTCGAGTATGCGTACTTCTTCATCAA G | Reverse primer no 4 for ICElm1 mobilization analysis |
| | **EMSA analysis** | |
| **Sif F_pET28a** | GAACATATGAAAAAAGCAATTTTAGATCGGATAGAAG | Forward primer, cloning of *sif* to pET28a. NdeI restriction enzyme site (underlined) |
| **Sif R_pET28a** | TGAGCTCAGTTTCTCCCTCTTAATCTTTCTAG | Reverse primer, cloning of *sif* to pET28a. SacI restriction enzyme site (underlined) |
| ***hfq*_F_EMSA** | CGCGAAAAGACAGTTAACGTGGT | Forward primer for EMSA DNA of *hfq* |
| ***hfq*_R_EMSA** | GCGACATTCTTCTGCGGGG | Reverse primer for for EMSA DNA of *hfq* |
| ***cadA*_F_EMSA** | CTGGTCTGTAAGGGTAGCAC | Forward primer for EMSA DNA of *cadA* |
| ***cadA*_R_EMSA** | TGCATCAGTTACACCCTCAAG | Reverse primer for for EMSA DNA of *cadA* |
| ***argC*_F_EMSA** | GTAGCGGAAACTGCGCAAGTG | Forward primer for EMSA DNA of *argC* |
| ***argC*_R_EMSA** | AGCGAATTAGCTCAAGTCCTC | Reverse primer for for EMSA DNA of *argC* |
| ***lap*_F_EMSA** | TGCTACTTACTCTTCTTCAATGCT | Forward primer for EMSA DNA of *lap* |
| ***lap*_R_EMSA** | TGGTTGAAAAGTCCTTTTGTAAAGT | Reverse primer for for EMSA DNA of *lap* |
| ***prfA*_F_EMSA** | CTGCGCGTTGATGACAAAGG | Forward primer for EMSA DNA of *prfA* |
| ***prfA*_R_EMSA** | GCGGTATGTTTCCACTGTCG | Reverse primer for for EMSA DNA of *prfA* |
| ***recA* F EMSA** | CCGTAAAACAAGGCTTTCAGT | Forward primer for EMSA DNA of *recA* |
| ***recA* R EMSA** | ATCCGCCAACTCCTAAAGCA | Reverse primer for for EMSA DNA of *recA* |
| ***lmo1097* F EMSA** | AGTTTGTCTATCAAAGCCCACAAG | Forward primer for EMSA DNA of *lmo1097* |
| ***lmo1097* R EMSA** | TCTTGTGCAAAAACTAATCAGACC | Reverse primer for for EMSA DNA of *lmo1097* |
| ***16sRNA* F EMSA** | GTGCATTAGCTAGTTGGTAG | Forward primer for EMSA DNA of *16sRNA* |
| ***16sRNA* R EMSA** | CAACAGTACTTTACGATCCG | Reverse primer for for EMSA DNA of *16sRNA* |
